# Nuclear stiffening in neoplastic cells aggregates T cell exhaustion via pFAK/SP1/IL-6 axis in colorectal cancer

**DOI:** 10.1101/2024.07.24.604770

**Authors:** Hao Kong, Xiangji Wu, Qingxin Yang, Qian Sun, Xin Yu, Yu Dai, Chunwei Wu, Dai Xie, Lang Chen, Panpan Ma, Siqi Dai, Chong Chen, Peng Mi, Xiaodong Feng, Yong Peng, Hubing Shi, Dunfang Zhang, Nengwen Ke, Dan Cao, Zongguang Zhou, Bing Hu, Ping Wang, Lu Chen, Yun-Hua Liu, Qiang Wei, Hong Jiang

## Abstract

Nuclear abnormalities such as nuclear deformation are hallmarks of many diseases, including cancer. Accumulating evidence suggests that the dense and mechanically stiff tumor microenvironment promotes nuclear deformation in cancer cells. However, little is known about how nuclear deformation in neoplastic cells regulates immune exhaustion in the tumor microenvironment. Here, we found that lamin A/C-mediated nuclear stiffening in neoplastic cells promotes the nuclear translocation of phosphorylated focal adhesion kinase (pFAK), which is strongly correlated with the heterogeneity and exhaustion of CD8^+^ T cells within the spatial context of the tumor microenvironment in human colorectal cancer. Mechanistically, we revealed that increased nuclear tension within tumor cells promotes pFAK nuclear translocation, where nuclear pFAK was found to regulate SP1/IL-6-mediated T-cell exhaustion and the transcription of proinflammatory cytokines/chemokines. Pharmacological inhibition or disruption of pFAK nuclear translocation enhanced antitumor immune responses and synergistically potentiated αPD-1 and αTIM-3 immunotherapy by increasing CD8^+^ T-cell cytotoxicity and restoring exhaustion in preclinical models of colorectal cancer. These findings highlight the pivotal role of nuclear tension-mediated pFAK translocation into the tumor cell nucleus in regulating CD8^+^ T-cell exhaustion, suggesting that pFAK is a promising target for advancing cancer immunotherapy.

## INTRODUCTION

The treatment of colorectal cancer (CRC) remains challenging despite advances in multimodal therapy, such as radiation, chemotherapy, and surgery^1,2^. While these approaches improve local disease control, the high distant recurrence rate of nearly 30% and substantial side effects highlight the need for novel therapeutic strategies^3,4^. Immunotherapy has revolutionized cancer treatment for several malignancies, such as melanoma, but has shown limited benefit in colorectal cancer, particularly in microsatellite stable (MSS) tumors^5,6^. Only a small portion of patients with high microsatellite instability (MSI-H) in colorectal cancer, estimated to be approximately 15%, are eligible for immunotherapy targeting PD-1, PD-L1, or CTLA-4^7,8^. Most colorectal cancers are microsatellite stable (MSS) and are not responsive to immunotherapies^9,10^. This disparity highlights the urgent need for innovative immunotherapeutic approaches tailored to colorectal cancers.

Studies have shown that a unique tumor microenvironment characterized by a fibrotic stroma, exhaustion of CD8^+^ tumor-infiltrating lymphocytes (TILs), and extensive infiltration by immunosuppressive cell populations contributes to the failure of therapies^11^. While biomechanical forces from the fibrotic niche promote nuclear deformation in cancer cells^12^, potentially by modulating nuclear tension and intracellular signaling^13^, their impact on T-cell function and tumor progression remains poorly understood. On the other hand, the heterogeneity and exhaustion of CD8^+^ T cells within the spatial context of the tumor microenvironment and their correlation with treatment response and patient survival are still largely unknown^14^. Critically, the molecular drivers of CD8^+^ T-cell exhaustion and immunotherapy resistance in CRC remain elusive^15^, representing significant challenges in cancer immunotherapy^16^. A better understanding of these mediators is needed to develop more effective immunotherapies for colorectal cancer and improve patient outcomes.

Nuclear deformation plays a critical role in modulating cellular functions, including cell migration and human disease pathogenesis^17^. Emerging evidence reveals that nuclear tension modulates the conformational state of nuclear pore complexes (NPCs), and the resulting nuclear deformations have a considerable effect on nucleocytosolic transport^18^. Studies have shown that mechanical signals from ECM rigidity are transmitted to the nucleus via LINC complexes^19^. These forces cause nuclear envelope stretching, likely opening nuclear pores and promoting nuclear transport^20^. Notably, the fibrotic, mechanically rigid tumor microenvironment induces pronounced nuclear deformation in cancer cells. How these biomechanical alterations in neoplastic cells regulate immune exhaustion in the tumor microenvironment is largely undefined.

Focal adhesion kinase (FAK) is a well-known nonreceptor protein tyrosine kinase that clusters at focal adhesion structures and plays a role in regulating cell adhesion, migration and survival. Our previous work and others have shown that targeting FAK in pancreatic cancer cells reduces the number of tumor-promoting macrophages and MDSCs and improves the response to anti-PD1/CTLA4 checkpoint therapy^21^. Recently, we reported that *PTK2* was predominantly expressed in epithelial cells, particularly in stem-like epithelial cells (**supporting Fig. 1a-e**). Interestingly, *PTK2* was overexpressed in tumors, especially in the MMRp subtype (**supporting Fig. 1f-g**), suggesting that FAK activation may play a critical role in driving T-cell exhaustion in the MMRp subtype of CRC.

In the present study, we performed spatial transcriptomics analysis and multiplex immunohistochemical staining to elucidate the spatial heterogeneity and functional characteristics of CD8^+^ TILs in human colorectal cancer. Our findings revealed that increased nuclear tension within the tumor microenvironment facilitates the nuclear translocation of pFAK and that the activation of tumor cell nuclear FAK is correlated with TIL heterogeneity and exhaustion within the tumor microenvironment. Using MC38 and CT26 mouse models of colorectal cancer, we demonstrated that pharmacological inhibition or genetic loss of FAK enhances antitumor immune responses by suppressing SP1/IL-6-mediated T-cell exhaustion. Importantly, targeting FAK with an inhibitor enhances the efficacy of αPD-1 and αTIM-3 immunotherapy by increasing the cytotoxicity of tumor-resident CD8^+^ T cells and reversing exhaustion states in preclinical models. These findings highlight FAK as a nuclear mechanosensor and support its inhibition as a promising strategy to improve immunotherapy outcomes in patients with colorectal cancer.

## RESULTS

### Nuclear stiffening promotes pFAK nuclear translocation and augments T-cell exhaustion in CRC

To investigate potential differences in nuclear tension across different cell types, we performed spatial transcriptomic analysis on human CRC tissues^22^. The tissues were annotated into five histologically distinct regions: tumor, epithelium, fibroblast, lamina propria, and smooth muscle. Among these regions, the tumor region presented markedly higher nuclear tension-related pathway scores (nuclear lamina) than the other regions did (**Fig. 1a**). Nuclear transport pathways displayed a similar spatial distribution and were strongly correlated with nuclear tension scores (R = 0.85, *P* < 2.2 × 10^−16^), indicating a tight association between elevated nuclear tension and enhanced nuclear transport activity in tumor cells (**Fig. 1a, b**). Lamin A/C constitute a critical structural component of the nuclear lamina and have been identified as predominant regulators of nuclear tension^23^. Immunohistochemical (IHC) analyses revealed significantly elevated lamin A/C protein expression in tumor regions compared with both adjacent normal tissues and nontumor areas (**Fig. 1c**), providing pathological evidence for the aberrantly increased nuclear tension characteristic of malignant cells and suggesting enhanced nuclear protein transport in tumor cells. Multiplex immunohistochemical (mIHC) staining revealed significantly increased nuclear translocation of pFAK in human CRC tissues than in adjacent normal tissues (**Fig. 1d**). Specifically, almost 80% of tumor cells presented positive nuclear pFAK staining, and tumor cells exhibiting high nuclear tension presented markedly greater pFAK nuclear accumulation than did their low-tension normal counterparts in normal tissue (**Fig. 1e**). In addition, we compared regions with high and low nuclear pFAK expression in the same CRC tissue and observed that high nuclear pFAK expression in tumor cells correlated with increased TIM-3 expression in adjacent T cells (**Fig. 1f, g**).

**Fig. 1.**
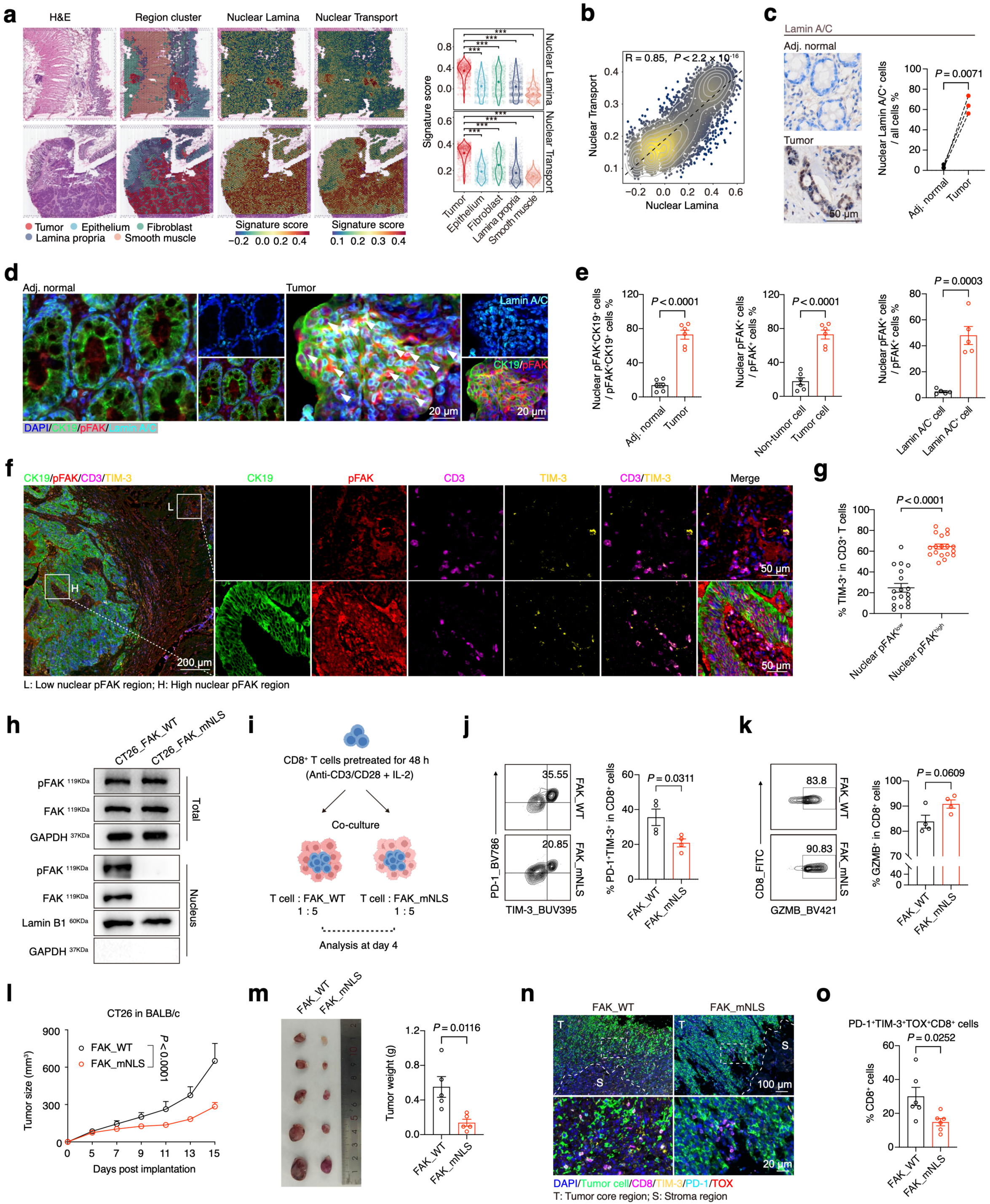
Nuclear stiffening enhances pFAK nuclear translocation and promotes T-cell exhaustion in the CRC TME. **a**, Spatial transcriptomics data illustrating the differential expression of the nuclear lamina and nuclear transport pathways across the tumor, epithelial, fibroblast, lamina propria and smooth muscle regions in CRC tumor tissue (left). Pathway activity scores were compared among these regions (right), with *P* values calculated via multiple two-sided unpaired t tests, \*\*\**P* < 0.0001. **b**, Spearman correlation between nuclear lamina and nuclear transport expression in CRC spatial transcriptomics data (R = 0.85, *P* < 2.2 × 10^−16^), with the *P* value calculated via a two-sided Spearman rank correlation test. **c**, Representative immunohistochemistry images and percentages of lamin A/C in human adjacent normal colon and CRC tumor tissues. The data are presented as the means ± SEMs (n = 3), as determined by two-tailed paired t tests. **d**, Representative mIHC images of pFAK distribution in tumor cells (lamin A/C high) and normal epithelial cells (lamin A/C low) stained with DAPI (blue), lamin A/C (cyan), pFAK (red) and CK19 (green). The arrows indicate tumor cells with pFAK nuclear localization. **e**, Comparisons of the percentage of nuclear pFAK^+^CK19^+^ cells between adjacent normal and tumor tissues (left) (n = 6 per group). Comparisons of the percentage of nuclear pFAK^+^ cells between nontumor cells and tumor cells in tumor tissue (middle) (n = 6 per group). Comparisons of the percentage of nuclear pFAK^+^ cells among lamin A/C^+^ cells and lamin A/C^−^ cells in tumor tissue (right) (n = 5 per group). The data are presented as the means ± SEMs, as determined by two-tailed unpaired t tests. **f** and **g**, Representative mIHC images (f) and quantification of TIM-3^+^CD3^+^ T cells in low-nuclear pFAK regions or high-nuclear pFAK regions (ROIs, n = 18 per group) (b) of human colorectal cancer sections (g). The data are presented as the means ± SEMs, as determined by two-tailed unpaired t tests. **h**, Western blot analysis of total and nuclear FAK/pFAK expression in CT26 FAK_WT and FAK_mNLS cell lysates. **i**, Schematic diagram illustrating the *in vitro* coculture system of CD8^+^ T cells with FAK_WT or FAK_mNLS tumor cells. **j**, Percentages of PD-1^+^TIM-3^+^CD8^+^ T cells among CD8^+^ T cells in coculture systems (n = 4 per group). The data are presented as the means ± SEMs, as determined by two-tailed unpaired t tests. **k**, Percentage of GZMB^+^CD8^+^ T cells among CD8^+^ T cells in the coculture system (n = 4 per group). The data are presented as the means ± SEMs, as determined by two-tailed unpaired t tests. **l**, Size of CT26 FAK_WT or FAK_mNLS subcutaneous tumors (n = 5 per group). The data are presented as the mean ± SEM, as determined by two-way ANOVA. **m**, Tumor image and weights of CT26 FAK_WT and FAK_mNLS subcutaneous tumors (n = 5 per group). The data are presented as the mean ± SEM, as determined by a two-tailed unpaired t test. **n**, Representative mIHC images of DAPI (blue), tumor cells (green), CD8 (pink), PD-1 (cyan) and TOX (red) in CT26 FAK_WT and FAK_mNLS tumors (left). Comparisons of the percentages of terminally exhausted T cells (PD-1^+^TIM-3^+^TOX^+^) among CD8^+^ T cells within tumor core regions from CT26 FAK_WT and FAK_mNLS tumors. The data are presented as the means ± s.e.m.s (ROIs, n = 6 per group), as determined by two-tailed unpaired t tests.

To investigate whether tumor-intrinsic nuclear pFAK regulates T-cell exhaustion, we first used CRISPR-Cas9 gene editing to knock out *Ptk2* expression in CT26 tumor cells (termed CT26_sg*Ptk2*) (**Extended Data Fig. 1a**) and found that FAK deletion had no significant effect on cell proliferation (**Extended Data Fig. 1b**). Next, we established CT26_FAK_WT and CT26_FAK_mNLS cell lines by reintroducing either wild-type FAK or a nuclear localization signal (NLS)-mutant FAK into CT26_sg*Ptk2* cells. Western blot analysis of total and nuclear fractions confirmed impaired nuclear localization of NLS-mutant FAK (**Fig. 1h**). T cells were subsequently pretreated with anti-CD3/CD28 antibodies and interleukin (IL)-2 for 24 h before being cocultured with tumor cells to induce T-cell exhaustion (**Fig. 1i and Extended Data Fig. 1c**). Activated CD8^+^ T cells cocultured with CT26_FAK_mNLS or CT26_sg*Ptk2* tumor cells presented reduced T-cell exhaustion, as evidenced by a lower percentage of PD-1^+^TIM-3^+^ cells and a greater percentage of GZMB^+^ cells. In contrast, pronounced T-cell exhaustion was observed when the cells were cocultured with CT26_sgNTC or CT26_FAK_WT tumor cells (**Fig. 1j, k and Extended Data Fig. 1d, e**), suggesting that tumor nuclear pFAK can promote CD8^+^ T-cell exhaustion. Next, we assessed the effects of the FAK-NLS mutation and FAK knockout on tumor growth *in vivo*. Strikingly, FAK knockout did not result in a reduction in tumor volume in nude mice (**Extended Data Fig. 1f**), whereas both FAK-NLS mutation and FAK knockout resulted in a significant delay in tumor progression and reduced tumor weight in immunocompetent mice (**Fig. 1l, m and Extended Data Fig. 1g**), suggesting an enhanced antitumor immune response following nuclear FAK inhibition. To further validate the impact of tumor cell FAK-NLS mutation and FAK knockout on CD8^+^ T cells *in vivo*, we investigated the spatial exhaustion status of CD8^+^ T cells via mIHC staining (**Fig. 1n and Extended Data Fig. 1h**). Analysis of tumor core-infiltrating CD8^+^ T cells revealed a significant reduction in the percentage of PD-1^+^TIM-3^+^TOX^+^CD8^+^ terminally exhausted T (Tex) cells in both FAK-NLS-mutant and FAK-knockout tumors compared with that in control tumors (**Fig. 1o and Extended Data Fig. 1i**). These findings strongly suggest that the nuclear translocation of tumor-intrinsic FAK is a critical driver of T-cell exhaustion.

### Lamin A/C-mediated high-nuclear-tension neoplastic cells drive pFAK nuclear translocation

To investigate the specificity of FAK nuclear translocation in CRC tumor cells, we cultured HT29 and HCT116 tumor cells, as well as normal colonic epithelial NCM460 cells, under both 3D and adherent conditions *in vitro*. Consistent with observations in CRC tissues, mIHC staining of cell spheres and IF staining of adherent cells revealed that nuclear pFAK was exclusively detected in HT29 and HCT116 cells but not in NCM460 cells (**Fig. 2a, b**). Notably, HT29 and HCT116 cells exhibited unusually smooth nuclear morphologies, which contrasts with the pronounced folds and wrinkles found in the nuclear morphology of NCM460 cells (**Fig. 2c**), suggesting that the nuclei of tumor cells may exist in a state of greater tension. To quantify this phenomenon, nuclear tension was evaluated using a nanoindenter (**Fig. 2d**). The results demonstrated that tension in the nuclear region of HT29 and HCT116 cells was significantly greater than that in NCM460 cells, which aligns with the observed differences in nuclear morphology (**Fig. 2d**).

**Fig. 2.**
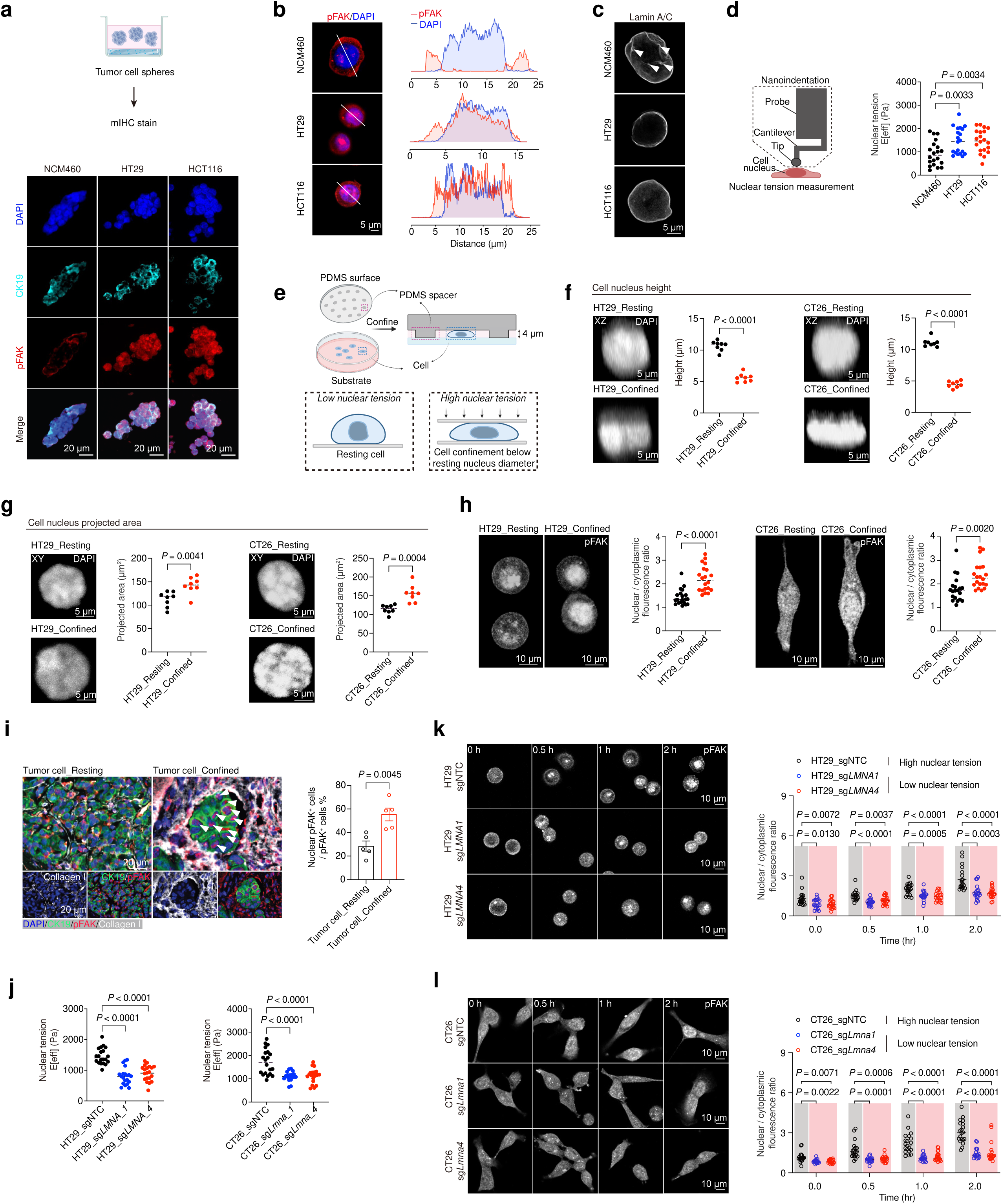
High nuclear tension in CRC tumor cells drives pFAK nuclear translocation. **a**, Representative merged images of 3D cell spheres stained with mIHC for DAPI (blue), CK19 (cyan) and pFAK (red). **b**, Representative immunofluorescence staining of pFAK in NCM460, HT29 and HCT116 cells (left) and fluorescence colocalization analysis of pFAK and DAPI in NCM460, HT29 and HCT116 cells (right). **c**, Images of the lamin A/C-labeled nuclear membrane in NCM460, HT29 and HCT116 cells. **d**, Schematic diagram of nanoindentation (left) and measurements of nuclear tension in NCM460, HT29 and HCT116 cells (right, n = 20 per group). The data are presented as the means ± s.e.m.s, as determined by one-way ANOVA. **e**, Schematic diagram of cell confinement. **f**, 3D xz views of the DAPI-stained nucleus. Measurements of nuclear height in resting and confined HT29/CT26 cells (n = 8 per group). The data are presented as the means ± SEMs, as determined by a two-tailed unpaired t test. **g**, 3D xy views of the DAPI-stained nucleus. Measurements of the nuclear projection area in resting and confined HT29/CT26 cells (n = 8 per group). The data are presented as the means ± SEMs, as determined by two-tailed unpaired t tests. **h**, Nuclear-to-cytoplasmic pFAK fluorescence ratio in resting and confined HT29/CT26 cells incubated with leptomycin B for 0.5 hours (n = 20 per group). The data are presented as the means ± SEMs, as determined by two-tailed unpaired t tests. **i**, Representative mIHC images of pFAK distribution in resting and confined tumor cells stained with DAPI (blue), CK19 (green), pFAK (red) and Collagen I (white) (n = 5 ROIs per group). The arrows indicate tumor cells with pFAK nuclear localization. The data are presented as the means ± SEMs, as determined by two-tailed unpaired t tests. **j**, Measurements of nuclear tension in HT29/CT26_sgNTC and sg*LMNA*/*Lmna* cells (n = 20 per group). The data are presented as the means ± s.e.m.s, as determined by one-way ANOVA. **k**, Representative immunofluorescence images and the nuclear-to-cytoplasmic pFAK fluorescence ratio in HT29_sgNTC and sg*LMNA* cells incubated with leptomycin B for 0, 0.5, 1, or 2 hours (n = 20 per group). The data are presented as the means ± s.e.m.s, as determined by one-way ANOVA. **l**, Representative immunofluorescence images and the nuclear-to-cytoplasmic pFAK fluorescence ratio in CT26_sgNTC and sg*LMNA* cells incubated with leptomycin B for 0, 0.5, 1, or 2 hours (n = 20 per group). The data are presented as the means ± s.e.m.s, as determined by one-way ANOVA.

On the basis of our findings, we aimed to investigate whether nuclear tension drives pFAK nuclear translocation. To address this, we first employed a cell confinement device to apply mechanical pressure to the nucleus (**Fig. 2e**). Using the tumor cell lines HT29 and CT26, we conducted a pressure experiment in which the height of individual nuclei was refined from an average resting diameter of approximately 10 μm to 4 μm (**Fig. 2e**). We characterized the nuclear morphology and deformation under 4 μm confinement and observed that while the nuclear height decreased (**Fig. 2f**), the projected surface area of the nucleus increased significantly (**Fig. 2g**), indicating a potential increase in nuclear tension. The cells were subsequently subjected to confinement for 0.5 hours in the presence of leptomycin B (a nuclear export inhibitor), followed by rapid fixation and staining for cellular pFAK. Our analysis revealed that the nuclear localization of pFAK was significantly greater in confined cells than in resting cells, a trend that was observed in both the HT29 and CT26 cell lines (**Fig. 2h**). *In vivo*, we compared tumor cells with high versus low nuclear pFAK localization via mIHC. The results indicated that cells exhibiting signs of mechanical extrusion, as evidenced by their encasement of abundant collagen, were associated with high nuclear pFAK localization (**Fig. 2i**). These findings suggest that increased nuclear tension promotes pFAK nuclear translocation.

We further tested whether pFAK translocation was affected by Lamin A/C-mediated nuclear tension. Lamin A/C is an important component of the nuclear skeleton that mediates the maintenance of nuclear pores and morphology. Previous studies have shown that lamin A/C expression is positively correlated with nuclear tension^23^, and our western blot results also revealed that lamin A/C expression in tumor cells (high nuclear tension) was greater than that in normal epithelial cells (low nuclear tension) (**Extended Data Fig. 2a**). We knocked down lamin A/C in both human (HT29 and HCT116) and mouse (CT26 and MC38) CRC tumor cells and found that lamin A/C-depleted cells displayed nuclear dysmorphia and increased nuclear folds, reflecting the relaxation of tension in the nuclear envelope (**Extended Data Fig. 2b-e**). To estimate this parameter, nuclear tension was examined in sg*LMNA*/*Lmna* cells and sgNTC cells, and as expected, the nuclear tension of sg*LMNA*/*Lmna* cells was significantly lower than that of sgNTC cells, both in human and mouse CRC tumor cells (**Fig. 2j and Extended Data Fig. 2b-e**). Then, pFAK nuclear import was measured by monitoring the nuclear accumulation of pFAK while blocking nuclear export with leptomycin B for 0 h, 0.5 h, 1 h, and 2 h. In both HT29 and CT26 cells, lamin A/C-deficient (low nuclear tension) cells presented significantly reduced nuclear import of pFAK compared with wild-type controls (high nuclear tension), indicating that impaired nuclear translocation of pFAK was caused by reduced nuclear tension in lamin A/C-deficient cells (**Fig. 2k, l**). Similar to the *in vitro* results, normal intestinal epithelial cells with low lamin A/C expression in adjacent normal tissues presented no expression of nuclear pFAK, whereas tumor cells with high lamin A/C expression in tumor tissues presented obvious nuclear localization of pFAK (**Fig. 1d and e**).

Taken together, our data suggest a novel mechanism by which tumor cells regulate the nuclear localization of pFAK through abnormally high nuclear tension. We propose that high collagen I expression in tumor tissues and high lamin A/C expression in tumor cells affect tumor nuclear tension, which controls pFAK nuclear translocation.

### Tumor nuclear pFAK promotes IL-6 secretion by regulating the transcription factor SP1

To elucidate the potential mechanism of nuclear pFAK-targeted genes, we first performed cleavage under targets and tagmentation (CUT&Tag) analysis in DMSO- or FAKi-treated CT26 cells. The results of western blot and immunofluorescence (IF) staining demonstrated that FAKi treatment significantly suppressed pFAK expression in CT26 cells, accompanied by a corresponding reduction in nuclear pFAK fluorescence (**Fig. 3a and Extended Data Fig. 3a**). Analysis of promoter regions (< 3 kb) with signals detected in the intersecting genes from DMSO vs. pFAKi 12 h, 24 h, and 12 h vs. 24 h CUT&Tag data from CT26 cells revealed that pFAK binds to promoters of 1,299 genes (**Fig. 3b, c**), and by ranking these genes, we identified the top-ranked transcription factor, SP1 (specificity protein 1) (**Fig. 3d**). We next analyzed the pFAK-binding motifs via the hybrid optimization of multiple energy resources (HOMER) motif discovery program 40^24^ (**Fig. 3e**) and found that the top-ranked motif was located in the pFAK binding site of the *Sp1* promoter region. Through DNA pull-down assays, we demonstrated that the pFAK protein specifically binds to the motif-containing sequence in the *Sp1* promoter (**Extended Data Fig. 3b**). Notably, FAKi treatment reduced the DNA-binding activity of pFAK to the *Sp1* locus (**Fig. 3f**). Validation via CUT&Tag-qPCR revealed a significant reduction in pFAK enrichment at the *Sp1* promoter and a corresponding decrease in *Sp1* mRNA expression following FAKi treatment compared with those in DMSO-treated control CT26 cells (**Fig. 3g, h**). Furthermore, we performed IF staining for pFAK and SP1 in FAK_WT and FAK_mNLS CT26 cells (**Fig. 3i, j**). The results revealed that SP1 was expressed mainly in the cell nucleus and that SP1 fluorescence was significantly lower in FAK_mNLS cells than in FAK_WT cells, as shown by fluorescence quantitative analysis (**Fig. 3j**). To validate the relationship between nuclear pFAK and SP1 in human CRC, we conducted mIHC staining on adjacent normal and tumor tissues (**Fig. 3k**). Overall, SP1 expression was significantly higher in tumor tissues than in adjacent normal tissues. Specifically, compared with normal cells with low nuclear tension (low lamin A/C expression), tumor cells with high nuclear tension (high lamin A/C expression) and distinct nuclear pFAK translocation exhibited significantly upregulated SP1 expression (**Fig. 3k**).

**Fig. 3.**
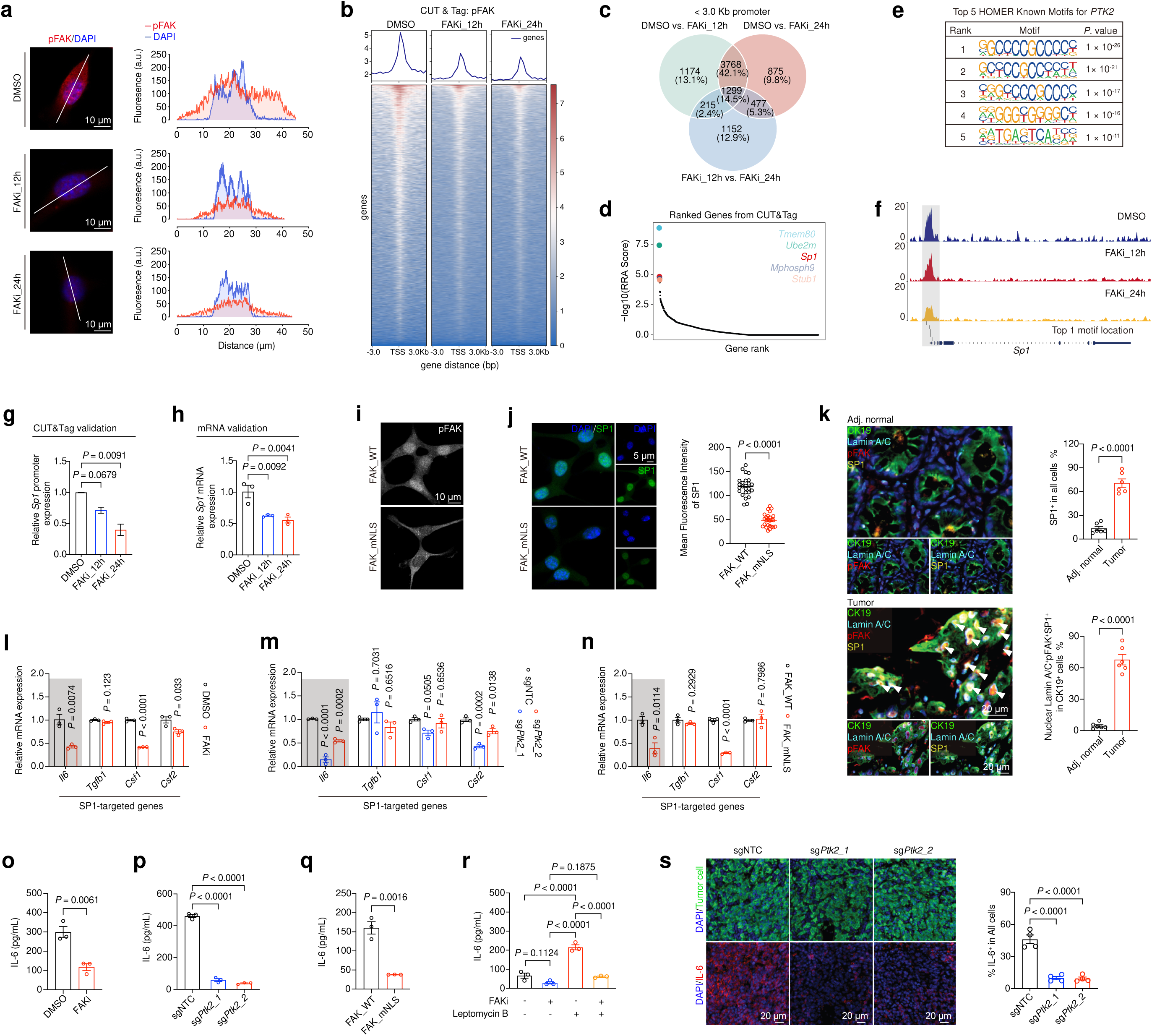
Tumor nuclear pFAK promotes IL6 secretion by regulating the transcription factor SP1. **a,** Representative immunofluorescence staining of pFAK in CT26 cells treated with DMSO or FAKi for 12 and 24 hours (left), fluorescence colocalization analysis of pFAK and DAPI in CT26 cells treated with DMSO or FAKi for 12 and 24 hours (right). **b**, Heatmap of CUT&Tag sequencing data showing pFAK binding sites in CT26 cells treated with DMSO or FAKi for 12 and 24 hours (n = 2 per group). **c**, Venn diagram showing the intersection of genes with promoter (< 3 kb) signals identified through MACS2 analysis across three comparisons: DMSO vs. FAKi 12 h, DMSO vs. FAKi 24 h, and FAKi 12 h vs. FAKi 24 h. **d**, Robust rank aggregation (RRA) ranking of the 1,299 CUT&Tag-enriched genes identified in Fig. 3c, with the top five ranked genes displayed. **e**, Top five known motifs of pFAK enriched by HOMER according to the peaks identified in the DMSO vs. FAKi comparison at 24 h. **f**, CUT&Tag sequencing tracks of the *Sp1* locus. Differentially accessible sites are highlighted with gray bars, and the top-ranked motif in the promoter region is indicated by a gray horizontal line. **g**, Real-time quantitative PCR (RT‒ qPCR) of the pFAK-binding promotor of *Sp1* in the CUT&Tag library constructed from DMSO- and FAKi (12 and 24 hours)-treated CT26 cells (n = 2 per group). The data are presented as the mean ± SEM, as determined by one-way ANOVA. **h**, Real-time quantitative PCR (RT‒qPCR) of *Sp1* transcript levels in CT26 cells treated with DMSO or FAKi for 12 or 24 h (n = 3 per group). The data are presented as the mean ± SEM, as determined by one-way ANOVA. **i**, Representative immunofluorescence staining of pFAK in CT26 FAK_WT and FAK_mNLS cells. **j**, Representative immunofluorescence staining of SP1 in CT26 FAK_WT and FAK_mNLS cells and mean fluorescence intensity analysis of SP1 in CT26 FAK_WT and FAK_mNLS cells (n = 25 per group). The data are presented as the mean ± SEM, as determined by a two-tailed unpaired t test. **k**, Representative mIHC images of SP1 expression in tumor cells (lamin A/C and nuclear pFAK high) and normal epithelial cells (lamin A/C and nuclear pFAK low) stained with DAPI (blue), CK19 (green), lamin A/C (cyan), pFAK (red) and SP1 (yellow). The arrows indicate the nuclear lamin A/C^+^pFAK^+^SP1^+^ tumor cells. Comparisons of the percentage of SP1^+^ cells in all cells (ROIs, n = 6 per group) and the percentage of nuclear lamin A/C^+^pFAK^+^SP1^+^ CK19^+^ cells (ROIs, n = 6 per group) between adjacent normal and tumor tissues. The data are presented as the means ± SEMs, as determined by two-tailed unpaired t tests. **l**, RT‒qPCR analysis of cytokine transcript levels in CT26 cells treated with DMSO or FAKi for 24 h (n = 3 per group). The data are presented as the mean ± SEM, as determined by a two-tailed unpaired t test. **m**, RT‒qPCR of cytokine transcript levels in CT26 sgNTC and sg*Ptk2* cells (n = 3 per group). The data are presented as the mean ± SEM, as determined by one-way ANOVA. **n**, RT‒qPCR analysis of cytokine transcript levels in CT26 FAK_WT and FAK_mNLS cells (n = 3 per group). The data are presented as the mean ± SEM, as determined by a two-tailed unpaired t test. **o-q**, IL-6 secretion measured by ELISA in supernatants from DMSO-treated, FAKi-treated (24 hours), sgNTC, sg*Ptk2*, FAK_WT, and FAK_mNLS CT26 cells stimulated with 20 ng/ml LPS for 6 hours *in vitro* (n = 3 per group). The data are presented as the mean ± SEM, as determined by one-way ANOVA or two-tailed unpaired t test. **r**, IL-6 secretion measured by ELISA in supernatants from FAKi (24 h)- and LMB (1 h)-treated CT26 cells stimulated with 20 ng/ml LPS for 6 hours *in vitro* (n = 3 per group). The data are presented as the mean ± SEM, as determined by one-way ANOVA. **s**, Representative mIHC images and percentages of IL-6^+^ cells in CT26 sgNTC and sg*Ptk2* tumors (n = 4 per group). The data are presented as the means ± SEMs, as determined by one-way ANOVA.

On the basis of the finding that pFAK regulates the transcription factor SP1, we subsequently validated the expression of SP1-targeted cytokine/chemokine genes^25^ and found that *Il6*, *Csf1*, and *Csf2* were consistently downregulated at the mRNA level following FAK inhibition in CT26 cells (**Fig. 3l**). Consistent with these findings, *Il6* and *Csf1* expression was also reduced in CT26 cells with FAK knockout or FAK_mNLS mutations (**Fig. 3m, n**). Using forward-phase protein arrays (FPPAs), which enable the quantification of 40 chemokines/cytokines in conditioned media from CT26 cells, we observed that FAK inhibition led to broad reprogramming of chemokines/cytokines, including the SP1 target IL-6 (**Extended Data Fig. 4a-c**). Next, we focused on IL-6 to investigate its role in T-cell regulation. Owing to the low basal secretion of IL-6 in CT26 cell culture media, we stimulated cells with lipopolysaccharide (LPS) and measured the IL-6 levels via ELISA to validate the FPPA results. ELISAs confirmed that FAK inhibition, FAK knockdown, and FAK_mNLS mutation significantly reduced IL-6 secretion compared with that in control cells, indicating that both nuclear FAK and FAK activation are critical for IL-6 regulation (**Fig. 3o-q**). To further demonstrate that nuclear pFAK regulates IL-6 secretion, we treated CT26 cells with leptomycin B to increase pFAK nuclear localization (**Extended Data Fig. 3c**). Increased nuclear localization of pFAK resulted in a significant increase in IL-6 secretion, which was fully reversed by FAKi treatment (**Fig. 3r**). To evaluate this relationship *in vivo*, we performed IL-6 staining in control and sg*Ptk2* tumors and found that IL-6 was nearly undetectable in FAK-deficient tumors (**Fig. 3 s**). These results suggest that the kinase activity of nuclear FAK is essential for the SP1-dependent regulation of IL-6 secretion.

### Tumor-intrinsic IL-6 promotes CD8^+^ T-cell exhaustion

To investigate the downstream effects of tumor cell-derived IL-6 on T-cell exhaustion, splenic CD8^+^ T cells were isolated and then exposed to IL-6 *in vitro* (**Fig. 4a**). The results revealed that IL-6 significantly promoted the expression of the exhaustion genes *Pdcd1* and *Havcr2* in T cells, whereas the expression of the effector gene *Ifng* was suppressed by IL-6 (**Extended Data Fig. 5 a, b**). We further confirmed that IL-6 promoted the expression of exhaustion markers, such as PD-1, TIM-3 and TOX, in CD8^+^ T cells and increased the proportion of exhausted CD8^+^ T cells (PD-1^+^TIM-3^+^TOX^+^CD8^+^ T cells) in a time-dependent manner (**Fig. 4b**). To identify the downstream mediators of IL-6-induced T-cell exhaustion, we examined the activation status of STAT3, a key downstream target of IL-6 signaling^26^. As expected, IL-6 promoted the expression and activation of STAT3 in a time-dependent manner (**Fig. 4c and Extended Data Fig. 5c, d**).

**Fig. 4.**
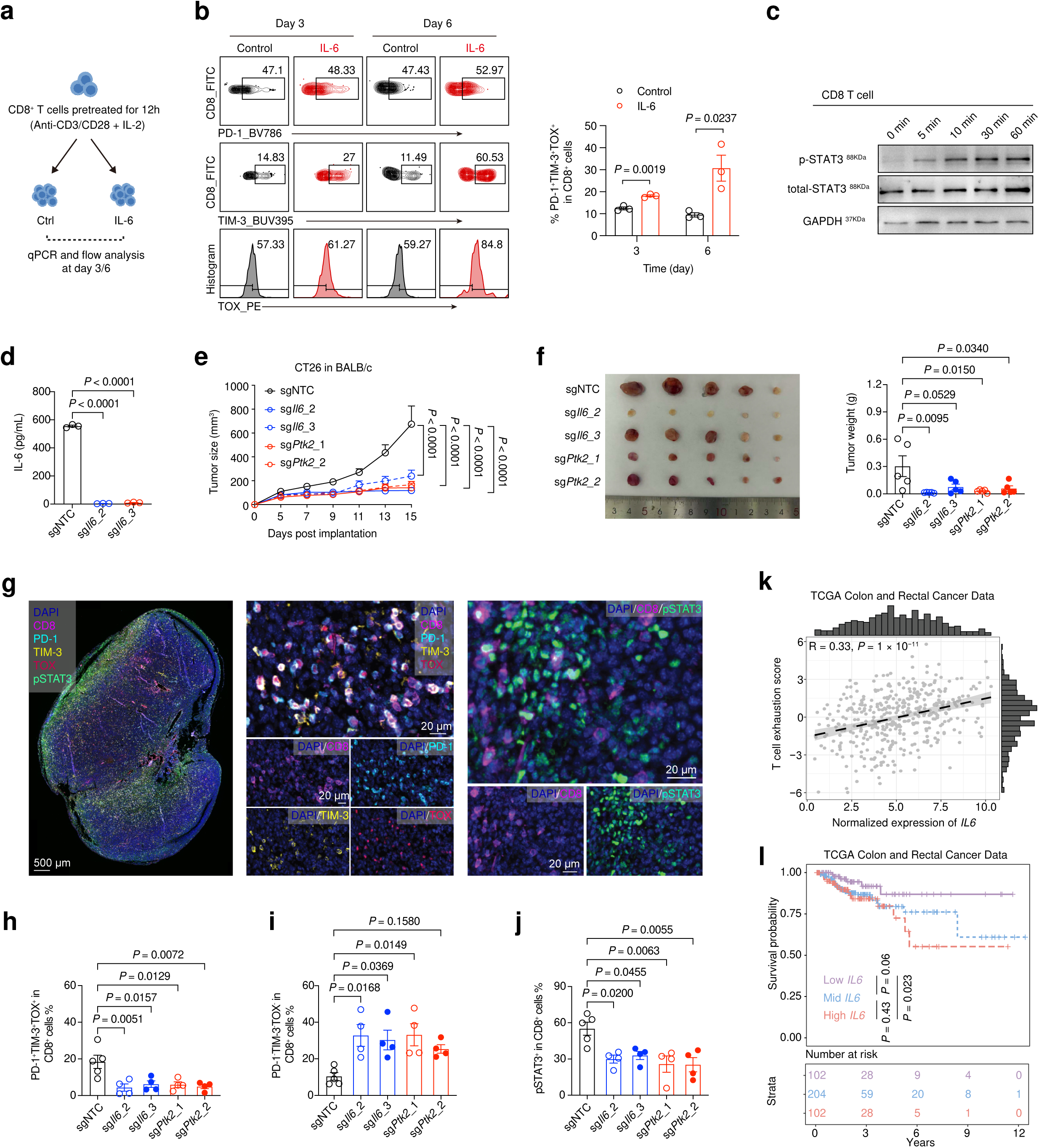
Nuclear pFAK-regulated IL-6 promotes CD8^+^ T-cell exhaustion. **a**, Schematic diagram of the *in vitro* culture model. **b**, Percentages of PD-1^+^TIM-3^+^TOX^+^CD8^+^ T cells among CD8^+^ T cells treated with or without IL-6 for 3 days or 6 days, as determined by flow cytometry (n = 3 per group). The data are presented as the means ± SEMs, as determined by two-tailed unpaired t tests. **c**, Western blot analysis of pSTAT3 and total STAT3 expression in CD8^+^ T cells treated with IL-6 over time. The time-dependent experiment was performed with a fixed IL-6 concentration of 10 ng/mL. **d**, IL-6 secretion in supernatants from CT26 sgNTC and sg*Il6* cells stimulated with 20 ng/ml LPS for 6 hours *in vitro* was measured by ELISA (n = 3 per group). The data are presented as the mean ± SEM, as determined by one-way ANOVA. **e**, Growth curves of CT26 sgNTC, sg*Ptk2* and sg*Il6* tumors in BALB/c mice (n = 5 per group). The data are presented as the means ± SEMs, as determined by two-way ANOVA. **f,** Tumor images and weights (n = 5 per group). The data are presented as the means ± SEMs, as determined by one-way ANOVA. **g**, Representative merged images of tumor sections stained with mIHC for DAPI (blue), CD8 (pink), PD-1 (cyan), TIM-3 (yellow), TOX (red) and pSTAT3 (green). **h**, Comparisons of the percentages of PD-1^+^TIM-3^+^TOX^+^CD8^+^ cells among CT26 sgNTC, sg*Ptk2* and sg*Il6* tumors (n = 4–5 per group). The data are presented as the means ± SEMs, as determined by one-way ANOVA. **i**, Comparisons of the percentages of PD-1^−^TIM-3^−^TOX^−^CD8^+^ cells from CT26 sgNTC, sg*Ptk2* and sg*Il6* tumors (n = 4–5 per group). The data are presented as the means ± SEMs, as determined by one-way ANOVA. **j**, Comparisons of the percentages of pSTAT3^+^CD8^+^ cells from CT26 sgNTC, sg*Ptk2* and sg*Il6* tumors (n = 4–5 per group). The data are presented as the means ± SEMs, as determined by one-way ANOVA. **k**, Spearman correlation between *IL6* expression and the T-cell exhaustion signature in TCGA-CRC bulk RNA-seq data (R = 0.33, *P* = 1 × 10^−11^), with the *P* value calculated via a two-sided Spearman rank correlation test. **l**, Kaplan–Meier survival analysis of TCGA-CRC patients stratified by *IL-6* expression quartiles, with *high IL-6 levels* ( ≥ 75^th^ percentile, red), *moderate IL-6 levels* (25th–75th percentile, blue), and *low IL-6 levels* (≤ 25th percentile, purple) included.

To validate the role of IL-6 in modulating CD8^+^ T-cell function *in vivo*, we generated IL-6-knockdown CT26 cell lines (**Fig. 4d**) and inoculated sg*Il6*, sg*Ptk2*, and sgNTC cells into BALB/c mice. Knocking down IL-6 and FAK in tumor cells markedly blunted tumor progression and led to almost complete regression of some tumors (**Fig. 4e, f**). We subsequently evaluated the exhaustion status of CD8^+^ T cells via mIHC (**Fig. 4g**). Compared with sgNTC tumors, sg*Il6* and sg*Ptk2* tumors presented a decreased percentage of PD-1^+^TIM-3^+^TOX^+^CD8^+^ terminal exhausted T cells and an increased percentage of PD-1^−^TIM-3^−^TOX^−^CD8^+^ functional T cells (**Fig. 4h, i**). Consistently, the loss of IL-6 and FAK in tumor cells led to a reduced proportion of pSTAT3^+^CD8^+^ T cells (**Fig. 4j**). Bioinformatic analysis of the Cancer Genome Atlas (TCGA) datasets revealed a positive correlation between IL-6 expression and the T-cell exhaustion score in patients with colorectal cancer (R = 0.33, *P* = 1 × 10⁻¹¹; **Fig. 4k**). Survival analysis of patients stratified by IL-6 expression levels revealed that the low-expression group exhibited significantly longer survival than did the high-expression group (**Fig. 4l**). Collectively, these findings suggest that tumor-intrinsic nuclear pFAK promotes IL-6 secretion through the regulation of the transcription factor SP1, thereby driving T-cell exhaustion in the TME.

### Pharmacological inhibition of FAK leads to stable disease progression by reprogramming the TME

To investigate the impact of FAK inhibition on tumor progression and alterations in the TME, we assessed the effects of the clinically available FAK inhibitor VS-4718 in murine colorectal tumor models, including MC38 (microsatellite instability, MSI) and CT26 (microsatellite stable, MSS) models. Compared with control mice, MC38 tumor-bearing mice treated with 50 mg/kg FAK inhibitor (FAKi) presented significantly reduced tumor growth and stable disease (**Fig. 5a**). This tumor-suppressive effect was consistently observed in the CT26 subcutaneous tumor model (**Fig. 5b**). IF staining of CT26 tumors revealed that nuclear pFAK and IL-6 expression was significantly inhibited after treatment with FAKi (**Fig. 5c, d**).

**Fig. 5.**
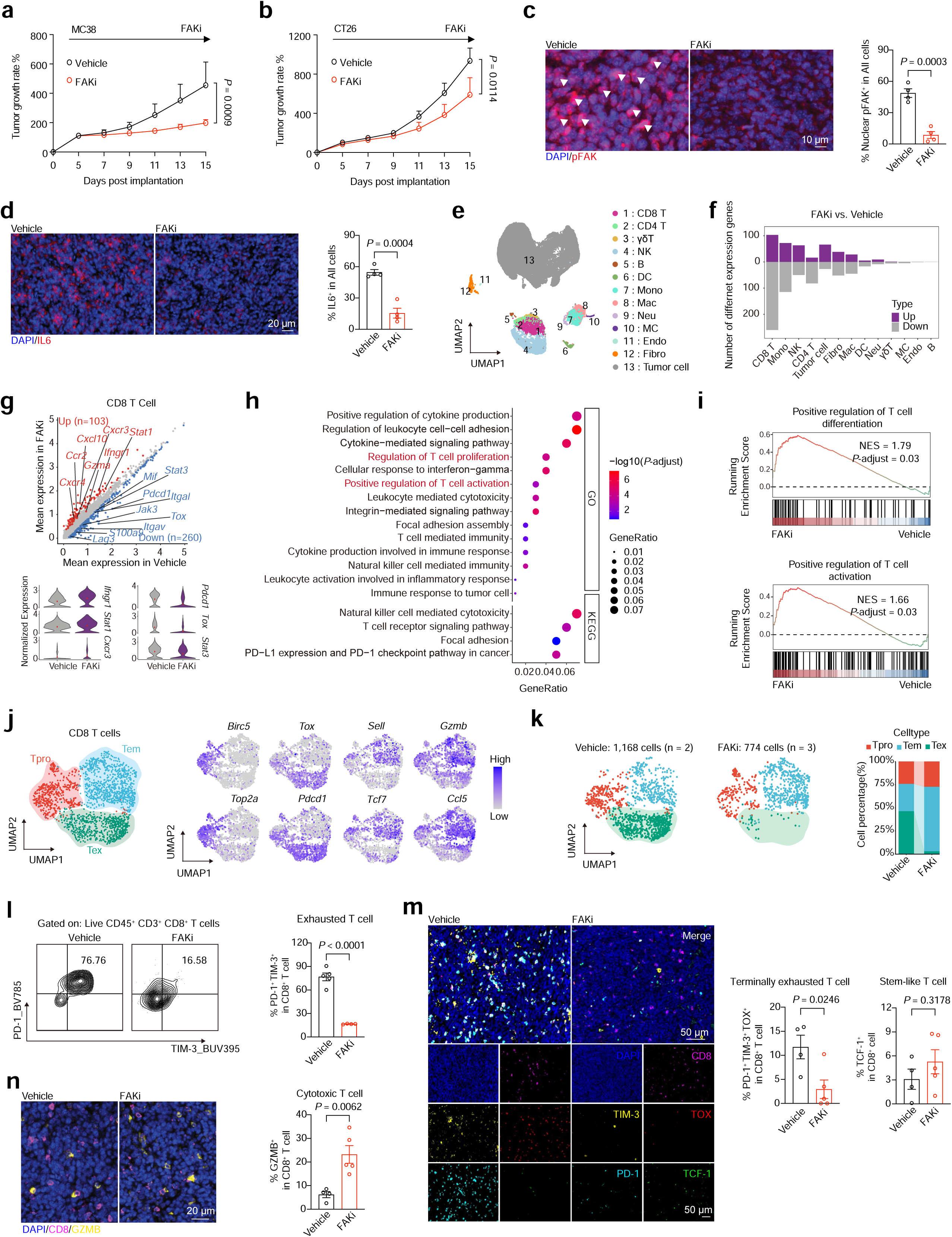
FAK inhibition leads to stable disease progression by reshaping the TIME. **a,** Growth rate of MC38 subcutaneous tumors treated with a FAK inhibitor (FAKi, VS-4718, 50 mg/kg) or vehicle (vehicle, n = 5; FAKi, n = 6). Vehicle or FAKi was administered intragastrically twice per day. The data are presented as the mean ± SEM, as determined by two-way ANOVA. **b,** Growth rate of subcutaneous CT26 tumors treated with a FAK inhibitor (FAKi, VS-4718, 50 mg/kg) or vehicle (n = 6 per group). Vehicle or FAKi was administered intragastrically twice per day. The data are presented as the means ± SEMs, as determined by two-way ANOVA. **c,** Representative mIHC images and percentages of nuclear pFAK^+^ cells in vehicle- or FAKi-treated tumors (n = 4 per group). The data are presented as the means ± SEMs, as determined by two-tailed unpaired t tests. **d**, Representative mIHC images and percentages of IL-6^+^ cells in vehicle- or FAKi-treated tumors. The data are presented as the means ± SEMs (n = 4 per group), as determined by two-tailed unpaired t tests. **e**, UMAP plot of 47,442 cells from vehicle (n = 2)- or FAKi (n = 3)-treated tumor tissues, colored according to the annotated cell type. **f**, Numbers of DEGs (upregulated in purple, downregulated in gray) in each cell type between the FAKi-treated group and the vehicle-treated group. Genes were identified on the basis of an adjusted *P* value < 0.05 and log₂-fold change > 0.25 via the Benjamini‒Hochberg (BH) method for multiple testing correction. **g,** Differentially expressed genes in CD8⁺ T cells between the FAKi-treated group and the vehicle group, as shown in Fig. 5f. The X-axis and Y-axis represent the average expression levels in the vehicle and FAKi groups, respectively. The upregulated genes are shown in red, and the downregulated genes are shown in blue (top). Selected representative genes were visualized via violin plots to illustrate expression differences between the two groups (bottom). **h**, GO and KEGG terms enriched for differentially expressed genes from Fig. 5g. *P* values were calculated via a hypergeometric test and adjusted for multiple comparisons via the Benjamini‒Hochberg (BH) method. The color scale represents the adjusted *P* values, and the symbol size reflects the number of enriched genes associated with each term. **i**, Gene set enrichment analysis of transcriptional signatures related to T-cell differentiation and activation in FAKi-treated vs. vehicle-treated CD8^+^ T cells. *P* values were calculated via a permutation test and adjusted for multiple comparisons via the Benjamini‒Hochberg (BH) method. **j**, UMAP plot of CD8^+^ T cells from all cells collected from the scRNA-seq data. CD8^+^ T cells were clustered into Tpro (red), Tem (blue), and Tex (green) subsets (left panel). Key markers for clustering are shown (right panel). **k**, Relative fraction of each cluster in Fig. 5j between the vehicle- and FAKi-treated groups. The left panel shows the UMAP plot, whereas the right panel presents the stacked bar chart. **l**, Percentages of exhausted (TIM-3^+^PD-1^+^) CD8^+^ T cells among CD8^+^ T cells determined by flow cytometry in CT26 tumors from vehicle-treated mice (n = 5) and FAKi-treated mice (n = 4 per group). The data are presented as the means ± SEMs, as determined by two-tailed unpaired t tests. **m,** Representative image and percentages of PD-1^+^TIM-3^+^TOX^+^CD8^+^ T cells and TCF-1^+^CD8^+^ T cells among CD8^+^ T cells in CT26 tumors from vehicle- or FAKi-treated mice (n = 4–5 per group). The data are presented as the means ± SEMs, as determined by two-tailed unpaired t tests. **n,** Representative image and percentage of GZMB^+^CD8^+^ T cells among CD8^+^ T cells in CT26 tumors from vehicle- or FAKi-treated mice (n = 4–5 per group). The data are presented as the means ± SEMs, as determined by two-tailed unpaired t tests.

Subsequent single-cell RNA sequencing (scRNA-seq) analysis of CT26 tumors enabled characterization of the transcriptional landscape of the tumor immune microenvironment (TIME). After quality control, we obtained transcriptomes from 47,442 cells and identified 13 main cell types, including various T/NK cell subsets, B cells, myeloid cells, endothelial cells, fibroblasts, and tumor cells (**Extended Data Fig. 6a-b**). The identified populations were subsequently visualized through dimensionality reduction via the uniform manifold approximation and projection (UMAP) algorithm (**Fig. 5e**). Notably, CD8^+^ T-cell gene changes were most significant after FAKi treatment compared with those in the control group (**Fig. 5f**). Compared with control tumors, FAKi-treated tumors presented upregulated expression of cytotoxicity-associated genes (*Stat1*, *Ifngr1*, *and Cxcr3*) and downregulated expression of exhaustion-related genes (*Stat3*, *Pdcd1*, *and Tox*) in CD8^+^ T cells (**Fig. 5g**). Gene Ontology (GO) enrichment analysis, Kyoto Encyclopedia of Genes and Genomes (KEGG) pathway analysis, and gene set enrichment analysis (GSEA) further revealed that FAKi treatment was associated with increased T-cell differentiation and activation pathways (**Fig. 5h, i**). Recluster analysis of 1,942 CD8^+^ T cells from both groups identified three subclusters, including proliferating CD8^+^ T cells (Tpro) expressing proliferation markers (*Bicr5* and *Top2a*), effector/memory CD8^+^ T cells (Tem) expressing effector/memory markers (*Gzmb*, *Ccl5*, *Sell* and *Tcf7*) and exhausted CD8^+^ T cells (Tex) expressing exhaustion markers (*Tox* and *Pdcd1*) (**Fig. 5j**). Notably, FAKi treatment dramatically reduced the percentage of the Tex population and increased the percentage of the Tem population (**Fig. 5k**).

Validation through mIHC and flow cytometric analysis confirmed that FAKi treatment significantly reduced the proportion of exhausted T cells (PD-1^+^TIM-3^+^CD8^+^ T cells and PD-1^+^TIM-3^+^TOX^+^CD8^+^ T cells) while increasing the percentage of stem-like CD8^+^ T cells expressing TCF-1 in CT26 tumors (**Fig. 5l, m and Extended Data Fig. 7**). Furthermore, FAKi treatment increased the proportion of GZMB^+^CD8^+^ T cells (effector CTLs) in CT26 tumors (**Fig. 5n**). Together, these findings suggest that FAK inhibition suppresses tumor progression by preventing T-cell exhaustion, maintaining a stem-like state, and enhancing T-cell effector functions, thereby reshaping the tumor immune microenvironment toward an antitumor response.

### FAK inhibitors potentiate immune checkpoint blockade (ICB) therapy *in vivo*

Given that FAK inhibition suppresses tumor growth by reprogramming the tumor microenvironment, we sought to further investigate its potential to increase the efficacy of immunotherapy. We found that FAKi synergized strikingly with immunotherapies when combined with low-dose 5-fluorouracil (20 mg/kg, i.p., ‘5-Fu^20^’) in both the CT26 (MSS) and MC38 (MSI) tumor models of human CRC, resulting in notable suppression of tumor growth compared with that in those treated with a FAK inhibitor or immunotherapy alone (**Fig. 6a-d**).

**Fig. 6.**
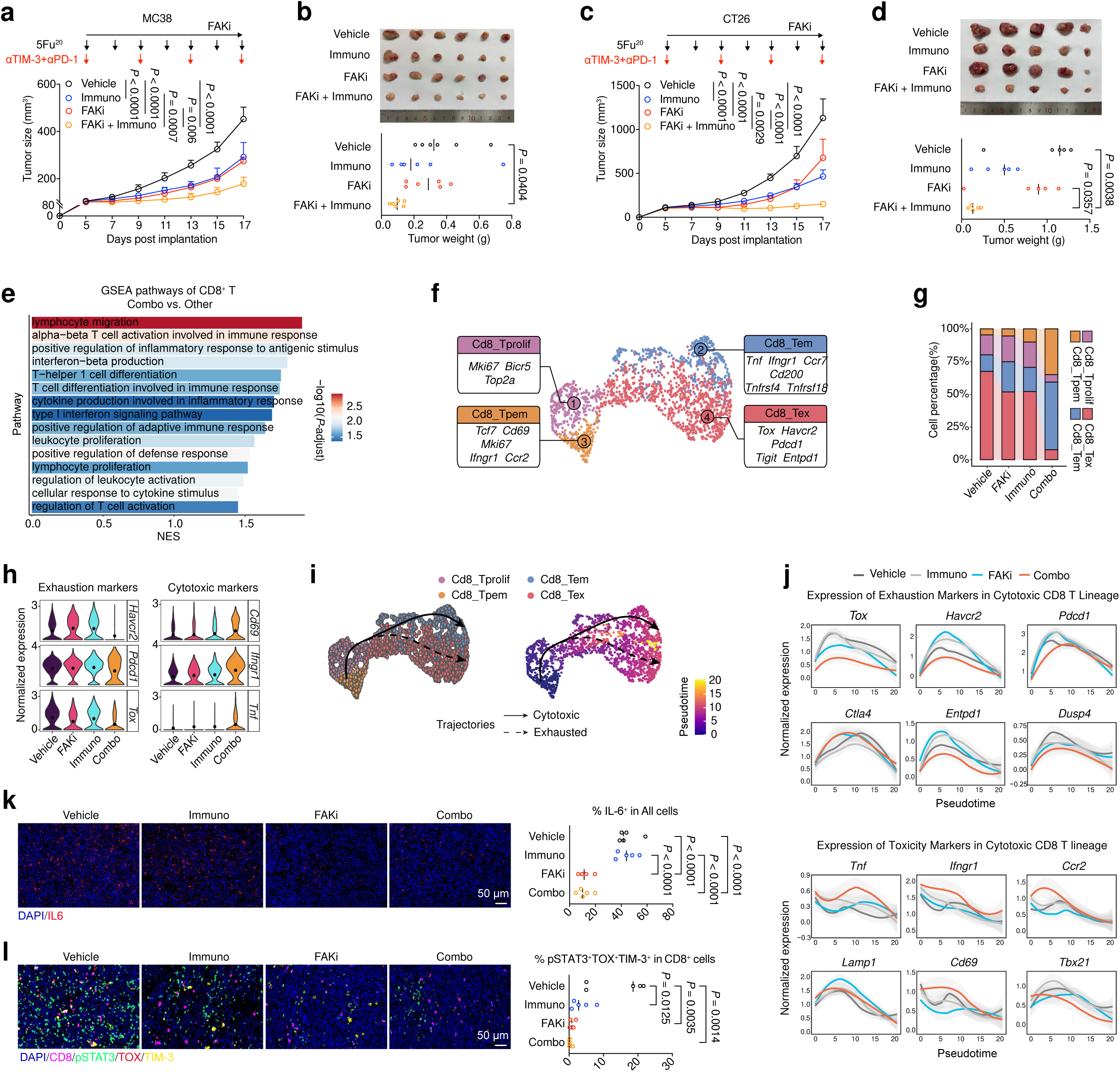
Inhibition of FAK improves the efficacy of immune checkpoint blockade (ICB) therapy *in vivo*. **a**, Growth curves of subcutaneous MC38 tumors treated with combination therapies (n = 6 per group). In the Immuno group, 5-Fu (20 mg/kg) combined with anti-TIM-3 (10 mg/kg) and anti-PD-1 (10 mg/kg) was used. Vehicle or FAKi was administered intragastrically twice per day. 5-Fu or vehicle was intraperitoneally administered once every two days. Isotype or anti-TIM-3 combined with anti-PD-1 was intraperitoneally administered once every 4 days. The data are presented as the means ± SEMs, as determined by two-way ANOVA. **b**, Tumor images and weights (n = 6 per group). The data are presented as the means ± SEMs, as determined by one-way ANOVA. **c,** Growth curves of subcutaneous CT26 tumors treated with combination therapies (n = 5 per group). In the Immuno group, 5-Fu (20 mg/kg) combined with anti-TIM-3 (10 mg/kg) and anti-PD-1 (10 mg/kg) was used. Vehicle or FAKi was administered intragastrically twice per day. 5-Fu or vehicle was intraperitoneally administered once every two days. Isotype or anti-TIM-3 combined with anti-PD-1 was intraperitoneally administered once every 4 days. The data are presented as the means ± SEMs, as determined by two-way ANOVA. **d**, Tumor images and weights (n = 5 per group). The data are presented as the means ± SEMs, as determined by one-way ANOVA. **e**, Gene set enrichment analysis of CD8^+^ T cells in the combination-treated group versus the other groups. *P* values were calculated via a permutation test and adjusted for multiple comparisons via the Benjamini‒Hochberg (BH) method. The color scale represents the – log10(adjusted *P* value). **f**, UMAP plot of CD8^+^ T cells from all the cells collected from the scRNA-seq data. CD8^+^ T cells were clustered into Tprolif (purple), Tpem (yellow), Tem (blue), and Tex (red) subsets. Key markers of the clustering process are highlighted in the map. **g,** Relative fraction of each cluster in Fig. 6f in the vehicle-, immune-, FAKi- or combination-treated groups. **h,** Violin map of the expression of exhaustion markers (*Havcr2*, *Pdcd1*, and *Tox*) and cytotoxic markers (*Cd69*, *Ifngr1* and *Tnf*) in the vehicle (purple), immunotherapy (red), FAKi (blue), and combination-treated (yellow) groups. **i,** Two trajectories (cytotoxic trajectory and exhausted trajectory) of CD8^+^ T cells based on Slingshot, and the CD8^+^ T-cell phenotype (left) and pseudotime (right) are displayed. The solid line represents the cytotoxic trajectory, whereas the dashed line represents the exhausted trajectory. **j,** Plot of exhaustion and cytotoxicity markers along the cytotoxic CD8^+^ T-cell lineage on the basis of pseudotime from Fig. 6i. The black line represents the vehicle group, the gray line represents the immunotherapy group, the blue line represents the FAKi group, and the red line represents the combination-treated group. **k,** Percentages of IL-6^+^ cells among all cells in CT26 tumors from vehicle-treated mice (n = 5), immuno-treated mice (n = 5), FAKi-treated mice (n = 4) and combination-treated mice (n = 5) determined by mIHC. The data are presented as the means ± SEMs, as determined by one-way ANOVA. **l,** Percentages of pSTAT3^+^TOX^+^TIM-3^+^CD8^+^ T cells among CD8^+^ T cells in CT26 tumors from vehicle-treated mice (n = 5), immuno-treated mice (n = 5), FAKi-treated mice (n = 4) and combination-treated mice (n = 5) determined by mIHC. The data are presented as the means ± SEMs, as determined by one-way ANOVA.

To decipher the underlying cellular and molecular mechanisms, we conducted scRNA-seq on tumor-infiltrating CD45^+^ immune cells (**Extended Data Fig. 8a**). After quality control, T-distributed stochastic neighbor embedding (t-SNE) analysis identified a total of 11 main cell types on the basis of known expression markers (**Extended Data Fig. 8b, c**). GSEA revealed that CD8⁺ T cells from combination-treated tumors were enriched in pathways related to T-cell functions, such as migration, proliferation, activation, and interferon-beta production (**Fig. 6e**). Reclustering analysis of CD8⁺ T cells revealed four subsets: precursor effector/memory T cells (Tpem) coexpressing stem-like (*Tcf7*, *Cd69*, *Mki67*) and effector/memory markers (*Ifngr1*, *Ccr2*); proliferating T cells (Tprolif) characterized by *Mki67*, *Bicr5*, and *Top2a*; effector/memory T cells (Tem) marked by *Tnf*, *Ifngr1*, and *Cd200*; and exhausted T cells (Tex) expressing *Tox*, *Havcr2*, and *Pdcd1* (**Fig. 6f and Extended Data Fig. 8d**). Notably, we observed significant expansion of Tpem clusters and Tem cell clusters in combination-treated tumors, whereas the proportions of Tex cells were dramatically lower in combination-treated tumors than in the other three groups (**Fig. 6g**). Furthermore, compared with those from the other three groups, CD8^+^ T cells from combination-treated tumors presented reduced expression levels of exhaustion-related genes (*Havcr2*, *Pdcd1* and *Tox*) and increased expression levels of cytotoxicity-related genes (*Cd69*, *Ifngr1* and *Tnf*) (**Fig. 6h**). Pseudotime trajectory analysis, in which Tpem cells were used as the root population, revealed two branches of differentiation via Tprolif cells: one branch toward Tem cells and the other branch toward Tex cells, giving rise to cytotoxic and exhausted CD8⁺ T-cell lineages, respectively (**Fig. 6i**). We observed that there were significantly lower levels of exhaustion marker genes in the cytotoxic CD8^+^ T lineage from the combination-treated tumors than in the other three groups. In contrast, the expression of cytotoxic marker genes was significantly greater in CD8^+^ T cells from the combination-treated group than in those from the other three groups (**Fig. 6j and Extended Data Fig. 8e**).

To validate the findings from the scRNA-seq data, we carried out mIHC to profile the CD8^+^ T-cell status in both CT26 tumors. We confirmed that there was a significant decrease in IL-6 protein expression in the combination-treated group as well as in the FAKi alone group (**Fig. 6k**). We also detected a significant reduction in pSTAT3^+^TOX^+^TIM-3^+^CD8^+^ T cells in the combination and FAKi groups (**Fig. 6l**).

Taken together, these findings demonstrate that inhibition of FAK enhances checkpoint therapy across CRC models by potentiating tumor-resident CD8^+^ T-cell functionality and cytotoxicity and reinvigorating exhausted states through suppression of the SP1‒IL-6 proinflammatory axis.

## DISCUSSION

This work demonstrated that the increased nuclear pFAK induced by nuclear stiffening in tumor cells is a key regulator of CD8^+^ T-cell exhaustion in CRC. Mechanistically, we elucidated that increased nuclear tension within tumor cells facilitates the nuclear translocation of pFAK. Once in the nucleus, pFAK orchestrates SP1/IL-6-mediated T-cell exhaustion and regulates the transcription of proinflammatory cytokines and chemokines. Importantly, our findings provide evidence that targeting pFAK with small molecule inhibitors enhances the efficacy of αPD-1 and αTIM-3 immunotherapies by increasing the degree of cytotoxicity among tumor-resident CD8^+^ T cells and reversing exhausted states in mouse models of CRC.

In solid tumors, including CRC, the infiltration of T cells into tumor beds has long been associated with favorable outcomes^27^. However, how spatial immune microenvironments influence T-cell exhaustion and immunotherapy resistance remains poorly understood^28^. Moreover, the mediators contributing to immune suppression and the limited response to immunotherapy in patients with colorectal cancer remain unexplored. Our previous work and others have shown that targeting FAK in pancreatic cancer cells reduces the number of tumor-promoting macrophages and MDSCs and improves the response to anti-PD1/CTLA4 checkpoint therapy^21^. Here, by integrating spatial transcriptomics and single-cell RNA sequencing, we identified pFAK nuclear translocation as a driver of immune suppression and T-cell exhaustion in human CRC. Genetic or pharmacological disruption of pFAK nuclear transport reprograms the tumor microenvironment, reversing terminal exhaustion (PD-1^+^TIM-3^+^CD8^+^ T cells) and potentiating antitumor immunity in preclinical models.

Recent advances have revealed a close relationship between nuclear mechanical properties and malignant tumor behavior. During metastasis, tumor cells need to cross a narrow extracellular matrix (ECM) or vascular endothelial space (<5 μm), whereas nuclei (usually >10 μm) must undergo significant deformation to pass through^29^. The groundbreaking work of Denais *et al*. revealed that tumor cell migration through narrow spaces can induce nuclear envelope rupture and subsequent DNA damage^12^. Shah *et al*. expanded this finding by demonstrating that nuclear deformation can trigger DNA damage by increasing replication stress, which may promote genomic instability and accelerate the tumor metastatic process^30^. These discoveries establish nuclear mechanical adaptability as a critical factor in cancer development. Consistent with our findings, Swift *et al*. reported that lamin A/C expression and phosphorylation status directly regulate nuclear rigidity^31^, with aberrant lamin A/C expression frequently observed in malignant tumors. According to clinical CRC data, patients with elevated lamin A/C expression exhibit nearly twice the mortality rate of lamin A/C-negative patients^32^. Furthermore, these nuclear mechanical properties also participate in a broader “ECM stiffness–cytoskeletal tension–nuclear deformation” feedback loop that activates mechanosensitive pathways, alters chromatin organization, and ultimately reinforces proinvasive gene programs. In our work, we demonstrated that nuclear stiffening in tumor cells can accelerate immune exhaustion in the tumor microenvironment by promoting FAK nuclear translocation, while disruption of pFAK nuclear translocation suppresses T-cell exhaustion, suggesting that nuclear mechanics might play a critical role in modulating immune suppression in colorectal cancer.

FAK, a nonreceptor tyrosine kinase localized at focal adhesions, regulates cell adhesion, migration, and survival. David Schlaepfer reported nuclear FAK translocation and its scaffolding role in cell survival^33^. Recently, Margaret Frame demonstrated that nuclear FAK is associated with chromatin and interacts with TFs to control evasion of antitumor immunity^34^. Consistent with these reports, we also observed that pFAK is located in the cell nucleus in the context of colorectal cancer. Importantly, our data suggest that increased nuclear tension, likely driven by the mechanically stressed tumor microenvironment, facilitates pFAK nuclear translocation, supporting a mechanoresponsive role for pFAK. We speculate that pFAK functions directly as a core transcriptional machinery. More importantly, we identified SP1 as a direct target of nuclear pFAK. We propose a model in which nuclear pFAK binds to the promoter of *Sp1*, which induces SP1 transcription and thereby leads to an increase in the production of IL-6, a direct target of Sp1^25^. Indeed, we demonstrated that IL-6 was a key contributor to T-cell exhaustion through the activation of STAT3 both *in vitro* and *in vivo*. Notably, FAK inhibition also expands NK cell populations, suggesting broader immunomodulatory effects beyond T-cell regulation, although the detailed underlying mechanisms warrant further exploration.

T-cell exhaustion, which is characterized by elevated expression of inhibitory receptors (PD-1, TIM-3, LAG-3, and CTLA-4), represents a critical barrier to effective antitumor immunity^11,35,36^. Targeting PD-1 with checkpoint inhibitors, such as nivolumab and pembrolizumab, has been shown to be effective in reversing T-cell exhaustion and restoring antitumor responses, leading to clinical benefits in some cancer patients^37,38^. However, while targeting PD-1 has shown promising results in some cancers, a significant proportion of patients still do not respond to this treatment^39,40^, highlighting the need for additional therapeutic targets and a better understanding of the complex mechanisms underlying T-cell exhaustion in cancer. T-cell immunoglobulin and mucin-domain-containing protein 3 (TIM-3) is considered an immune checkpoint molecule similar to PD-1 and CTLA-4^41,42^. Studies have highlighted the role of TIM-3 expression on CD8^+^ T cells in relation to disease stage in human colorectal cancer (CRC), as well as the potential benefits of TIM-3 blockade in enhancing antitumor responses^43,44^. These outcomes highlight the complexity of cancer immunotherapy and the need for multifaceted treatment approaches. Emerging evidence suggests that the long-term persistence of TCF1^+^ stem-like T cells is important for achieving durable responses to immunotherapies, such as checkpoint inhibitors^45,46^. In this work, we demonstrated that the nuclear translocation of pFAK contributes to T-cell exhaustion in human colorectal cancers. Genetic or pharmacological disruption of pFAK nuclear transport reprograms the tumor microenvironment, specifically reducing the number of terminally exhausted T cells while enhancing antitumor immunity in preclinical models.

FAK overexpression and hyperactivation are associated with tumor progression, metastasis, and resistance to therapy in various cancers. As a result, FAK inhibitors are being investigated in multiple clinical trials, both as monotherapies and in combination with chemotherapy, immunotherapy, or targeted therapies^47,48^. Our previous work demonstrated that a combination of FAK inhibitors and anti-PD-1/CTLA4 antibodies reduced the number of immunosuppressive cells and improved overall survival in mouse models of human pancreatic cancer^21^. Consistent with previous reports^34^, here, we found that targeted inhibition of FAK dramatically improved the efficacy of anti-PD1/TIM-3 checkpoint therapy in both MSS and MSI mouse models of human CRC as a result of reinvigorating exhausted states of tumor-infiltrating lymphocytes and potentiating effector T-cell cytotoxicity. FAK inhibitors represent a promising therapeutic approach in oncology, particularly in combination with immunotherapy and targeted agents. Ongoing clinical trials are aiming to validate their role in overcoming treatment resistance and improving outcomes in aggressive cancers (NCT06072781, NCT02428270, NCT01849744, NCT06166836 and NCT05580445). The results of these studies will determine their potential for future clinical application and provide promising evidence that targeting the FAK pathway may represent a promising strategy for cancer immunotherapy.

In summary, our study established nuclear pFAK as a mechanoresponsive transcriptional regulator that drives CD8^+^ T-cell exhaustion through the SP1‒IL-6 axis in CRC. Notably, our data suggest that increased nuclear tension within tumor cells facilitates pFAK nuclear translocation, thereby linking tumor mechanical stress to immune dysfunction. Together, these insights provide a strong rationale for therapeutically targeting FAK to overcome immune exhaustion and improve the efficacy of immune checkpoint blockade in colorectal cancer and a framework for developing spatially and mechanically informed immunotherapeutic strategies.

## METHODS

### Human tumor specimen collection

Human colorectal cancer specimens were collected from patients with histologically confirmed colorectal cancer who underwent curative surgical resection at the Department of Gastrointestinal Surgery, West China Hospital, Sichuan University, China. This project was approved by the West China Hospital Medical Committee (2020--374).

### Multiplex immunohistochemistry (mIHC)

Multiplex IHC staining was performed using an Opal 7-color kit (Akoya Bioscience, US). Briefly, the sections were dewaxed with xylene for 20 minutes. Then, ethanol was used for rehydration. Microwave treatment was performed for antigen retrieval with antigen retrieval buffer. Next, all the sections were cooled at room temperature for 30 minutes. Endogenous peroxidase activity was blocked via Antibody Diluent/Block (Akoya Bioscience, US) for 10 minutes at room temperature. The slides were then incubated for 1 hour at room temperature with the primary antibody (details of the antibodies are described in **Supplemental Table 1**), for 20 minutes at 37°C with the secondary reagents, and for 10 minutes at room temperature with Opal working buffer. The above procedures were repeated for other antibodies, and the antibodies were removed by microwave treatment before another round of staining was performed. Nuclear staining was performed via incubation with DAPI (Akoya Bioscience, US) for 5 minutes at room temperature. Slides were scanned at 20× magnification via the Vectra Polaris system (Akoya Biosciences, US). Whole-tissue scans were analyzed with Qupath software (https://qupath.github.io) via area quantification and a cytonuclear module^49^.

### Statistics and reproducibility

All the statistical analyses were performed via Prism 10 (GraphPad). The data are presented as the mean ± SEM, and two-tailed Student’s t test, one-way ANOVA, or two-way ANOVA was used as indicated in the figure legends. The exact value of n (number of biological or experimental replicates) can be found in the figure legends, and the exact P values are indicated in the graphed data. A P value < 0.05 was considered to indicate statistical significance. Most experiments were carried out at least two or three times, and the findings of all the key experiments were reliably reproduced.

## Authors’ contributions

HJ, QW, YHL and PW designed the research; HK, PW and HJ wrote the paper; NWK and HK collected the clinical samples; HK, QXY and Lang C analyzed the clinical sample data; QXY, HK and PPM performed the scRNA-seq data analysis; QS and HK performed the cell mechanics assays; XJW and SQD performed the CRISPR knockdown of FAK; XJW and HK generated the *in vivo* mouse models; CWW and DX helped with the drug treatments; HK, XY and YD performed the mIHC and analyzed the data; ZGZ, DC and BH supervised the clinical research design; YHL, Lu C and HBS supervised the scRNA-seq data analysis; XDF, CC and PM supervised the *in vivo* studies; PW, DFZ and YP supervised the T-cell coculture experiments.

## Acknowledgments

The authors thank all members of the Hong Jiang laboratory (West China Hospital, Sichuan University) for their insightful comments regarding this work. The authors thank the Laboratory of Digestive Surgery, West China Hospital, for clinical sample preparation and quality control (QC) validation. The authors thank Xiaogang Wang for providing the HALO image analysis platform. This work was supported by the following funding: Natural Science Foundation of China grants #82073158 (Hong Jiang), #82072615 (Yun-Hua Liu), #82273274 (Yun-Hua Liu) and # T2222020 (Qiang Wei); the Leading Goose R&D Program of Zhejiang (2024c03142 to Yun-Hua Liu) and 1.3.5 project for disciplines of excellence from West China Hospital of Sichuan University (ZYYC24005, to Hong Jiang).

