## Supplementary figures for "Nuclear stiffening in neoplastic cells aggregates T cell exhaustion via pFAK/SP1/IL-6 axis in colorectal cancer"

### METHODS

#### Immunohistochemical staining

All involved tissue samples were processed into paraffin. Paraffin-embedded tissues were cut into 5-μm sections and subjected to immunohistochemistry. The slides were immersed in antigen retrieval buffer (Gene Tech, China) at a 1:500 dilution in distilled water. Antigen retrieval was then performed in a microwave oven for ~20 min. Then, 2% BSA (BioFroxx, Germany) was used for blocking for 10 min, and the primary antibody (details of the antibodies are described in **Supplemental Table 1**) was diluted in PBS containing 1% BSA and applied overnight. A secondary antibody (Thermo Fisher Scientific, US) was added after the slides were washed for 30 min. After washing, DAB solution (Gene Tech, China) was added for coloring.

#### Cell culture and treatment

The human cell lines HT29, HCT116 and NCM460 were provided by Dr. Hubing Shi (Sichuan University) and cultured under standard conditions specified by the manufacturer. CT26 and MC38 cells were obtained from the ATCC and cultured under standard conditions specified by the manufacturer. The cell lines were maintained in a humidified atmosphere containing 5% $CO_2$ at

37° C and were regularly tested and verified to be mycoplasma negative via a mycoplasma detection kit (HUABIO, China).

For the leptomycin B treatment, the cells were incubated with 370 nM leptomycin B (Beyotime, 1:1000 dilution from a 370 μM stock in ethyl alcohol to cell medium) or with a 1:1000 dilution of ethyl alcohol as a control.

FAKi treatment: Cells were incubated with 1 μM VS-4718 (Selleck, US) or with DMSO as a control.

**Confinement of cell populations**

To generate confined cells for population-based or biochemical studies, cell confinement was achieved via a custom-designed device[1]. This device comprises a polydimethylsiloxane (PDMS, SYLGARD 184, US) suction cup that presses a confining coverslip equipped with PDMS microspacers (micropillars) onto a cell-populated culture substrate. The height of the micropillars (4 μm) defines the spatial confinement height between the coverslip and the substrate. The molds for the PDMS microspacers were fabricated via standard photolithography techniques.

**Nanoindentation of cell nuclei**

Mechanical testing was conducted via a Piuma Nanoindenter (Optics11 Life, The Netherlands). A spherical probe with a radius of 10 μm and a cantilever stiffness of 0.5 N/m was utilized. The cells were cultured in 3.5 cm diameter Petri dishes. Prior to measurement, the medium was replaced with PBS. The indentation protocol consisted of a loading phase of 2 s at an indentation depth of

3000 nm, held for one second, and then an unloading phase of 2 s. All individual indentation values were calculated via Piuma Data viewer version 2.2 (Piuma; Optics11, The Netherlands).

**Immunofluorescence staining**

Immunofluorescence staining was performed according to standard procedures. Briefly, the cells were fixed with 4% paraformaldehyde for 10 min, washed 3 times with PBS, permeabilized with 0.1% Triton X-100 in PBS for 5 min, incubated with primary antibodies overnight, and incubated with secondary antibodies for 1 h at room temperature. 4',6-Diamidino-2-phenylindole (DAPI) was added with the secondary antibodies. Fluorescence images were collected with a 60x oil immersion objective on a confocal microscope (Zeiss LSM 980, Carl Zeiss, Germany). Images were captured using identical exposure times for each cell line. Rabbit anti-pFAK and rabbit anti-lamin A/C antibodies were used as the primary antibodies (details of the antibodies are described in **Supplemental Table 3)**, and the secondary antibody was an AF555-labeled goat anti-rabbit IgG (H + L) antibody (Invitrogen, USA). The fluorescence was imaged via a laser confocal microscope (Leica-STELLARIS, Germany) and quantified via ImageJ.

**3D cell sphere culture**

To prepare polyHEMA-coated plates, 1 mL of polyHEMA solution (20 mg/mL in ethanol) was added to each well of a 6-well culture plate. The plate was gently swirled to ensure even coating across the well surfaces and allowed to dry overnight under a sterile hood. After drying, the coated plates were washed twice with sterile phosphate-buffered saline (PBS) to remove any residual ethanol. For spheroid formation, the cells were suspended in culture medium at the desired density and pipetted into the PolyHEMA-coated wells. The plate was gently swirled to ensure the even distribution of cells and then placed in a 37°C incubator with 5% $CO_2$. The cells were allowed to aggregate and form spheroids over 72 hours, depending on the cell type and experimental conditions. After spheroid formation, the spheroids were gently pipetted from the wells and transferred to a tube for fixation and embedding.

**3D cell sphere fixation and embedding**

The cell spheroids were fixed by submerging them in 4% paraformaldehyde (PFA) and incubating them at room temperature for 30 minutes. After fixation, the spheroids were washed three times with PBS. The fixed spheroids were centrifuged, and the excess PBS was removed. Then, prewarmed 2% agarose (37°C) was added to the spheroids, which were mixed gently to ensure even distribution. The spheroid-agarose mixture was quickly transferred to a microcentrifuge tube and allowed to solidify at 4°C for 10 minutes. The fixed cell spheroids were subjected to mIHC staining.

**Image analysis**

The slides were visualized via the Vectra Polaris system (Akoya Biosciences, US), and a multispectral image of the whole slide was scanned via a 20× objective lens. Multispectral image unmixing was performed via QuPath software (version 3.0)[2]. Briefly, the DAPI-positive cells were identified by the “cell detection” command, and each single channel intensity threshold was selected via “Object classification.” We determined the number of positive cells with the “Load classifier” and counted the areal proportions by dividing the channel-positive cell counts by all the cells detected as DAPI positive. Spatial distance is calculated as “distance to centroid distances 2D”. All other multispectral image unmixing methods were performed via HALO software. Image analysis algorithms were built via the Indica Labs High-Plex FL v2.0 module to perform DAPI-based nuclear segmentation and detect fluorescence-positive cells by setting a dye cytoplasm-positive threshold. The spatial analysis module was used for proximity analysis, infiltration analysis and density analysis.

**Cell proliferation**

For CT26_sgNTC and CT26_sg*Ptk2* cells, 2000 cells per well were seeded into 96-well plates in DMEM supplemented with 10% FBS. After 3 days and 6 days, CCK-8 solution (Life-ilab, China) was added to each well, and the plates were then incubated at 37°C. After 2 h, the absorbance at 450 nm was measured via a microplate reader (Thermo Fisher Scientific, US).

**Western blot**

Total proteins were extracted via RIPA lysis and extraction buffer (Thermo Fisher Scientific, US), and nuclear proteins were extracted via nuclear protein and cytoplasmic protein extraction kits (Beyotime, China). The protein concentration was assessed via a BCA protein assay kit (Life-ilab, China). Proteins were loaded into each lane of a polyacrylamide gel. After electrophoretic separation, the proteins were transferred to a PVDF membrane (Sigma–Aldrich, Germany). The membranes were blocked for 1 h in Tris-buffered saline containing 0.05% Tween 20 (TBS) and 5% nonfat dry milk. Then, the cells were incubated with a primary antibody (the details of the antibodies used are described in **Supplemental Table 1**) at 4°C overnight. The membranes were washed 3 times with TBS for 5 minutes each and then incubated with a secondary antibody (Thermo Fisher Scientific, US). An enhanced chemiluminescence (ECL) detection system (Tanon 5500, China) was used to image specific blots.

**Generation of stable cell lines**

To generate knockdown cell lines, guide RNAs (gRNAs) targeting the exons of the *Ptk2*, *Il-6*, *Lmna* and *LMNA* genes were designed via online tools available at https://www.atum.bio/eCommerce/cas9/input and https://www.genscript.com/gRNA-detail. The synthesized gRNAs were annealed and cloned and inserted into the BsmBI restriction sites of the V2TC plasmid, which encodes Cas9 and was generously provided by Dr. Chong Chen (Sichuan University). These recombinant plasmids were used as transfer vectors for lentiviral packaging. The gRNA sequences used were as follows:

| Guide RNA Names | Sequences |
|---|---|
| ms_*Ptk2*_gRNA1 | GTGCACCTCCTCCGATCGC |
| ms_*Ptk2*_gRNA2 | TCGAGTACTAAGACTCACC |
| ms_*Lmna*_gRNA1 | ACGCACGCGATCGATGTACA |
| ms_*Lmna*_gRNA4 | AGAGGTGTCCGGCATCAAGG |
| ms_*Il6*_gRNA2 | TGCAGAGAGGAACTTCATAG |
| ms_*Il6*_gRNA3 | CACCAGCATCAGTCCCAAGA |
| hu_*LMNA*_gRNA1 | GGCGAGCTGCATGATCTGCG |
| hu_*LMNA*_gRNA4 | TCGCTGGAAACGGAGAACGC |

For lentivirus production, human embryonic kidney 293T cells were transfected with the packaging plasmids pMD2. G, psPAX2, and the transfer plasmid via the calcium phosphate transfection method. In a 6-well plate, the following amounts of plasmids were used: 1 µg of pMD2.

G, 2 μg of psPAX2, and 4 μg of the transfer plasmid. The culture supernatants were collected at 36 and 48 hours posttransfection and filtered through a 0.45 μm membrane to isolate the lentiviral particles. Target cells were infected with lentivirus expressing both gRNA and Cas9, followed by selection with an appropriate concentration of puromycin for one week. The knockout efficiency of the target genes was validated via a T7 endonuclease assay and Western blot analysis.

To establish stable FAK-WT and FAK-mNLS CT26 cell lines, we initially amplified the native FAK-WT sequence from CT26 cellular cDNA and constructed recombinant lentiviral vectors through homologous recombination-mediated Cas9 replacement in the pLenti-Cas9-Zeocin backbone (Beyotime, China). Employing a multiround PCR-based mutagenesis approach, we engineered nuclear localization signal (NLS)-deficient FAK variants by introducing six critical point mutations (R177A/R178A/K190A/K191A/K216A/K218A, reported by Alan Serrels, *et al.*) into wild-type FAK (FAK-WT) via precision site-directed mutagenesis[3]. This combinatorial mutation strategy completely disrupted NLS functionality while preserving cytoplasmic retention, with no detectable alterations in basal protein expression profiles.

The NLS-mutated FAK construct was subsequently cloned and inserted into the same lentiviral expression system for parallel vector production. Following lentiviral particle packaging, we stably reconstituted FAK-WT or FAK-mNLS in FAK-knockout CT26 cells through optimized viral transduction protocols. The transduced populations were rigorously selected with 800 μg/mL Zeocin for 7 consecutive days to establish monoclonal cell lines. Successful genomic integration and mutant protein expression were systematically validated through western blotting and immunofluorescence localization analyses.

**Isolation and activation of CD8$^+$ T cells**

The spleens of C57BL/6 mice were dissected and ground with a syringe plunger, followed by filtration with a 40 μm cell filter and treatment with 1x RBC lysis buffer (Invitrogen, USA). The cells were collected by centrifugation at 1500 rpm for 5 min, and the naïve T cells were harvested via a MojoSort™ mouse CD8$^+$ T-cell isolation kit (#480008, Biolegend, US) following the manufacturer's instructions. For CD8$^+$ T-cell activation, 24-well flat-bottom plates were coated with 1.5 μg/mL anti-CD3 (#BE0001, BioXcell, US) and anti-CD28 (#BE0015, BioXcell, US) antibodies and incubated at 4°C overnight. The wells were subsequently washed with PBS, after which 5 × $10^5$ T cells were seeded and cultured at 37°C with 5% $CO_2$ in RPMI-1640 medium (HyClone, US) supplemented with 10% heat-inactivated fetal calf serum (ZETA LIFE, US), 1% penicillin and streptomycin (Gibco, US), 1% nonessential amino acids (Gibco, US), 1% sodium pyruvate (Gibco, US), 2% HEPES (Gibco, US), and 10 ng/mL IL-2 (#AF-212-12, PeproTech, US) for 24 h.

In an IL-6-induced Stat3 phosphorylation T-cell assay, T cells were activated in the presence or absence of recombinant mouse IL-6 (5, 10, 50, 100 ng/mL, #AF-216-16, PeproTech, US) for 0, 5, 10, 30, or 60 min, and p-Stat3 expression was assessed via western blot analysis.

In the IL-6-induced exhaustion of T cells assay, T cells were activated in the presence or absence of recombinant mouse IL-6 (10 ng/mL) for 3 days or 6 days, whereas exhaustion markers were assessed by qPCR and flow cytometric analysis.

**Tumor-CD8$^+$ T-cell coculture system**

Unless otherwise stated, CT26_sgNTC, CT26_sg*Ptk2*, CT26_FAK_wt and CT26_FAK_mNLS cells were seeded at a 5 : 1 ratio (tumor cells:$CD8^+$ T cells) into a 24-well plate. Activated T cells ($2 \times 10^5$ cells/well) were added to plates containing tumor cells. After approximately 4 days, all the cells were collected and analyzed via flow cytometry.

**Enzyme-linked immunosorbent assay (ELISA)**

ELISAs were carried out with a mouse IL-6 ELISA kit (#VAL604G, Novusbio, US) according to the manufacturer's instructions. After cell adhesion, the cells were starved for 3 h and subjected to LPS stimulation. After 6 h, the supernatant was collected for ELISA detection.

**Animal models**

The CT26 cells and MC38 cells were maintained at 37°C with 5% $CO_2$ in RPMI-1640 medium (HyClone, US) supplemented with 10% heat-inactivated fetal calf serum (ZETA LIFE, US), penicillin (Gibco, US), and streptomycin (Gibco, US). In all the animal experiments, the mycoplasma detection results for all the cell lines were negative. Six- to eight-week-old female C57BL/6, BALB/C and BALB/C nude mice were used for the subcutaneous tumor mouse models. Briefly, $2.5 \times 10^5$ CT26 cells and $5 \times 10^5$ MC38 cells in 50 µL of Matrigel (Corning, US) were injected subcutaneously into each mouse's right back. Tumor volume (length × (width$^2$)/2) was assessed by caliper measurements every other day, and cohorts of mice were randomized into different treatment groups according to tumor volume. All animal studies were approved by the West China Hospital Animal Ethics Committee (2020361A).

**Inhibitors, chemicals and neutralizing antibodies**

For the animal experiments, 50 mg/kg of VS-4718 (Selleck, US) was dissolved in vehicle (0.5% carboxymethyl cellulose (Sigma Aldrich, Germany) and 0.1% Tween-80 (Sigma Aldrich, Germany) in sterile water) and was administered orally by gavage every 12 hours. 5-Fu (#HY90006, MCE, USA) was dissolved in PBS and administered by intraperitoneal (i.p.) injection every 2 days at 20 mg/kg. Monoclonal anti-TIM-3 antibodies (#BE0115, BioXCell, US) and anti-PD-1 antibodies (#BE0146, BioXCell, US) were given by intraperitoneal (i.p.) injection every 4 days at 10 mg/kg. Isotype controls (#BE0089, BioXCell, US) were used at the same concentration in control mice.

**CUT&Tag**

The CUT&Tag assay was conducted following the manufacturer's protocol (#HD101, Vazyme, China). Briefly, Concanavalin A-coated magnetic beads were incubated with DMSO-treated CT26 or FAKi-treated CT26 cells at room temperature for 10 min, and anti-pFAK antibody (1:50, #700255, Invitrogen, US) was added and rotated at 4°C for 12 h. The samples were washed 3 times and then incubated with a secondary antibody (goat anti-rabbit IgG H&L, #AB206-01-AA, Vazyme, China) at 37°C for 30 min. After washing, the samples were incubated with Hyperactive pA/G-Transposon Pro and fragmented with $MgCl_2$. The fragmented DNA was extracted from the samples and amplified via PCR. Libraries were constructed via the TruePrep Index Kit V2 for Illumina (#TD202, Vazyme, China) and sequenced on an Illumina NovaSeq platform, and 150-bp paired-end reads were generated.

For CUT&Tag-qPCR, DNA samples were obtained according to the Hyperactive Universal CUT&Tag Assay Kit (#TD904, Vazyme, China), in which the target DNA fragments with DNA spike-in as a standard reference fragmented by pA/G-Tn5 were specifically recovered for qPCR to quantify the abundance of the target gene.

**Flow cytometry**

Single-cell suspensions of the tumors were obtained by incubating minced tissues with 3 mg/mL collagenase A (Roche, Germany) and 1 mg/mL DNase I (Roche, Germany) in RPMI 1640 medium at 37°C for 30 min. The resulting cell suspensions were passed through a 40 μM cell filter and treated with 1× RBC lysis buffer (Invitrogen, US). Finally, the suspension was washed with PBS, and the cells were counted for flow cytometric analyses. First, the cells were incubated with Fixable Viability Stain 700 to gate viable cells. The cells were subsequently washed with flow cytometry staining buffer (PBS containing 2% BSA), and the Fc receptor (FcR) was blocked with TruStain FcX™ (anti-mouse CD16/32) antibody (BioLegend, US). The cells were incubated with cell surface antibodies for 30 min at 4°C. The cells were then washed twice with flow cytometry staining buffer. For intracellular staining, the cells were cultured in fixation buffer at 4°C for 45 min (eBioscience, US). The cells were then washed twice with permeabilization buffer (eBioscience, US). The cells were incubated with the appropriate intracellular antibodies for an additional 30 min at 4°C. The details of the antibodies used are described in **Supplemental Table 2**. The cells were then washed twice with permeabilization buffer and resuspended in flow cytometry staining buffer.

**RT–qPCR**

RNA was prepared according to the TRIzol reagent (Invitrogen, US) protocol. After the generation of complementary DNA, qPCR with reverse transcription was performed as described previously, and all expression levels were normalized to those of an internal housekeeping gene (*Gapdh*). The primers used are listed in **Supplemental Table 3**.

**Protein arrays**

CT26 cells were stimulated for 12 h with FAKi (1 μM) or DMSO (Sigma Aldrich, Germany). The cell supernatants were then collected after centrifugation. Cytokines in the cell supernatants were then analyzed with the Proteome Profiler Mouse Cytokine Array Kit A (#ARY006, R&D Systems, US). The blots were visualized with chemiluminescence via a detection system (Tanon 5500, China) and quantified via ImageJ.

**Isolation of $CD45^+$ cells for scRNA-seq**

Single-cell suspensions of the tumors were obtained by incubating minced tissues with 3 mg/mL collagenase A (Roche, Germany) and 1 mg/mL DNase I (Roche, Germany) in RPMI 1640 medium at 37°C for 30 min. The resulting cell suspensions were passed through a 40 μM cell filter and treated with 1× RBC lysis buffer (Invitrogen, USA). Finally, the suspension was washed with PBS and counted for isolation. The Fc receptor (FcR) in the cells was subsequently blocked with a TruStain FcX™ (anti-mouse CD16/32) antibody (#101319, BioLegend, US). After blocking, the cells were incubated with a PE-conjugated anti-mouse CD45 antibody (#103106, BioLegend, US) for 20 min at 4°C. After washing, anti-PE MicroBeads (#130-048-801, Miltenyi, Germany) were added, and the mixture was mixed well and refrigerated for 15 minutes. After washing, the cell suspension was applied to the column, and labeled $CD45^+$ cells were collected for analysis.

**Bioinformatics**

**Cut&Tag data preprocessing**

The raw Cut-Tag reads were trimmed and filtered for quality via Cutadapt (v.4.6)[4]. Paired-end reads were aligned against mm10 genomes and spike-in sequences from *E. coli* genomes via Bowtie2 (v.2.5.1)[5] with the parameters --local --very-sensitive--local --no-mixed --no-discordant --phred33 -I 10 -X 700 -p 15. The aligned reads were sorted, and nonuniquely mapped reads were marked via Picard (v.3.0.0) (http://broadinstitute.github.io/picard/). PCR duplicates and reads of low mapping quality (MAPQ < 2) were filtered via SAMtools (v.1.18)[6].

We then identified peaks via MACS2 (v.2.2.7.1)[7], employing parameters with preprocessed bamfiles as follows: -f BAMPE -g mm -B -p 0.01. Peaks were annotated via the R package ChIPseeker (v.1.24.0)[8]. The enrichment of gene ontology terms in sets of peaks was calculated via ClusterProfiler (v.4.9.0.002)[9]. Motif analysis was performed via the default parameters of Homer (v.5.1)[10,] and a motif search was conducted via the R package MEME (v.1.13.1).

**Peak heatmaps and genome coverage plots**

To visualize CUT&Tag data coverage across the genome within protein-coding genes, read counts were normalized to the corresponding total reads aligned to the spike-in genome (*E. coli*). Normalized bigwigs were generated with the bamCoverage function from deepTools (v3.5.4)[11]. The heatmaps were generated with the indicated regions (TSSs ± 3,000 bp) with the computeMatrix function followed by the plotHeatmap functions of deepTools. Candidate gene coverage plots were generated via pyGenomeTracks (v3.8)[12].

**Single-cell transcriptomic analysis**

**Library preparation and sequencing**

The scRNA-seq libraries described in this study were generated via the DNBelab C4 system. Initially, a single-cell suspension was processed through several stages, including droplet encapsulation, emulsification, and fragmentation. This was followed by the collection of mRNA capture beads, reverse transcription, cDNA amplification, and subsequent purification. An indexed sequencing library was then assembled according to the manufacturer's protocol. The concentration of the sequencing library was determined via a Qubit ssDNA Assay Kit (Thermo Fisher, US). Sequencing was performed on the DIPSEQ T1 platform at the National Gene Bank (CNGB, BGI-SHENZHEN, China). For data processing, the raw FASTQ files were analyzed via DNBelab_C4scRNA (v1.0.1) software (available at https://github.com/MGI-tech-bioinformatics/DNBelab_C_Series_scRNA-analysissoftware) and converted into a format compatible with Cell Ranger[13].

**Alignment and quantification**

The sequencing data were processed via STAR software (v2.5.3)[5] with default parameters and mapped to the mm10 reference genome. The aligned reads were subsequently screened to identify valid cell barcodes and unique molecular identifiers (UMIs), which facilitated the creation of a gene–cell matrix for further analysis.

**Quantity of scRNA-seq data**

The gene–cell matrix was loaded by the Read10X function of the R package Seurat (v4.1.1)[14]. Genes expressed in fewer than 3 cells were identified. Additionally, the cells were filtered on the basis of four standards: UMIs, the number of genes, the percentage of mitochondrial (Mt) genes, and the percentage of ribosomal protein large/small (Rpl/s) genes. The percentage of Mt/Rpl/s genes was calculated via the percentage feature set function. These filtering standards vary across different datasets and depend on the data characteristics. For the FAK inhibitor experiments, cells were removed if they had UMIs less than 1100 or greater than 60000, fewer than 700 or greater than 7500, a Mt gene percentage greater than 15%, or an Rpl/s gene percentage greater than 30%. Similarly, in the therapeutic experiments, cells were excluded if they had UMIs less than 1000 or greater than 17000, number of genes less than 900 or greater than 4700, Mt gene percentage greater than 10%, or Rpl/s percentage greater than 30%. Additionally, only $CD45^+$ cells were retained in the therapeutic group samples. Following quality control, the analysis included 47,442 single cells from FAKi experiments and 75,631 from therapeutic experiments for downstream analysis.

**Dimensionality reduction and cell type classification**

After quality control of the cells and genes, dimensionality reduction and unsupervised clustering were performed via the R package Seurat (v4.1.1). To address batch effects, each sample was individually processed via the NormalizeData and FindVariableGenes functions. Subsequently, batch integration was achieved through the FindIntegrationAnchors and IntegrateData functions, setting the dims parameter to 50. This integrated dataset was then scaled for principal component analysis, utilizing the first 20 principal components to construct a shared nearest neighbor (SNN) network. The Louvain algorithm was employed to identify cell clusters, with resolution settings adjusted according to the specific characteristics of each dataset. Visualization of these clusters in two-dimensional space was accomplished via either UMAP or t-SNE, implemented via the RunUMAP or RunTSNE functions, respectively. Markers for each cluster were detected via the FindAllMarkers function in Seurat with the following parameters: min.pct = 0.1, logfc.threshold = 0.25, pseudocount.use = 0.1, only.pos = T. To annotate each cluster as a specific cell type, we selected some classic markers of immune cells, epithelial cells and fibroblasts. The cell types were annotated via a violin or dot diagram.

**Single-cell level differentially expressed gene (DEG) analysis**

Differential gene expression analysis was conducted via the FindMarkers function of the Seurat (v4.1.1) R package. For this analysis, the cell populations of interest were specified as ident.1 and ident.2. We calculated the fold changes in the mean gene expression levels between these populations. Only genes that exhibited a Bonferroni-adjusted *P* value of less than 0.05 and an absolute log2-fold change greater than 0.25 were selected for subsequent analysis.

**Slingshot pseudotime trajectory inference**

We used Slingshot (v1.6.0)[15] to construct cell trajectories and investigate cell-state transitions in $CD8^+$ T cells in therapeutic experiments. CD8 T cells were annotated as CD8_Tpem (*Tcf7*, *Cd69*, *Mki67*, *Ifngr1*, *Ccr2*), CD8_Tprolif (*Mki67*, *Bicr5*, *Top2a*), CD8_Tem (*Tnf*, *Ifngr1*, *Ccr7*, *Cd200*, *Tnfrsf4*, *Tnfrsf18*), or CD8_Tex (*Tox*, *Havcr2*, *Pdcd1*, *Tigit*, *Entpd1*). We selected CD8_Tem cells as the root state when the trajectories and pseudotime were calculated via the slingshot function. The analysis revealed two distinct lineages: cytotoxic and exhausted trajectories.

**Public Bulk RNA-seq, scRNA-seq, and spatial transcriptomics data analysis**

To evaluate whether FAK activation influences the TME, we performed an integrated single-cell analysis of 100 previously published scRNA-seq datasets (GEO: GSE178341). To examine the spatial differences in the nuclear lamina gene set, which was obtained from MSigDB (v.2023.2)[16], between tumor and adjacent normal tissues, we obtained spatial transcriptomics data from CRC patients via scCRLM[17]. The signature scores were computed via the GSVA (v.1.36.3) package[18]. Additionally, to assess the relationships among IL-6 expression, T-cell exhaustion, and patient survival, we obtained bulk RNA-seq data from TCGA-CRC via UCSC Xena.

**Exhaustion signature score calculation**

The exhausted signature consisted of eight well-established exhaustion-related genes: *PDCD1*, *HAVCR2*, *TOX*, *LAG3*, *CTLA4*, and *TIGIT*. For the bulk RNA-seq data, module scores for the exhaustion signature were computed via the gsva function from GSVA (v.1.36.3). We then calculated Spearman's correlation coefficients between IL-6 expression and the exhausted signature score for each sample to assess their associations.

**Survival analysis**

To investigate the relationship between the exhaustion signature and patient prognosis, we stratified patients into low-*IL6*, moderate-*IL6*, and high-*IL6* groups on the basis of their *IL6* expression levels. The top 25% (≥75th percentile) were classified as high-*IL6*, the bottom 25% (≤25th percentile) as low-*IL6*, and the remaining samples as middle-*IL6*. K–M survival curves were generated via the survfit function of the survival R package (v3.2-13) to compare prognostic differences between these groups.

**GO enrichment**

The enrichGO and enrichKEGG R packages ClusterProfiler (v.4.9.0.002) were used to perform Gene Ontology (GO) and Kyoto Encyclopedia of Genes and Genomes (KEGG) analyses via the org.Mm.eg.db (v.3.18.0) package. The results were visualized via the dotplot function of the R package enrichplot (v.1.20.0, https://github.com/YuLab-SMU/enrichplot).

### Gene Set Enrichment Analysis

Gene set enrichment analysis (GSEA) was performed via the clusterProfiler (v.4.9.0.002) package. Genes were initially sorted on the basis of their log2-fold change. The enrichment was then performed via the gseGO function with the org.Mm.eg.db (v3.18.0) package. Visualization was achieved with the GseaVis (v.0.0.5, https://github.com/junjunlab/GseaVis) package.

### Data and Code Availability

All the raw mouse sequencing data have been deposited in the CNCB (https://ngdc.cncb.ac.cn/) BIG subproject under the following accession numbers: PRJCA035685 (FAKi treatment single-cell data), PRJCA027119 (combination treatment single-cell data), and PRJCA036987 (CUT&Tag data). All sequencing data are publicly available as of the date of publication.

All software and R packages used are publicly available or previously reported. The analysis pipeline followed official guidelines and is detailed in the Methods section. The code is available from the corresponding authors upon request.

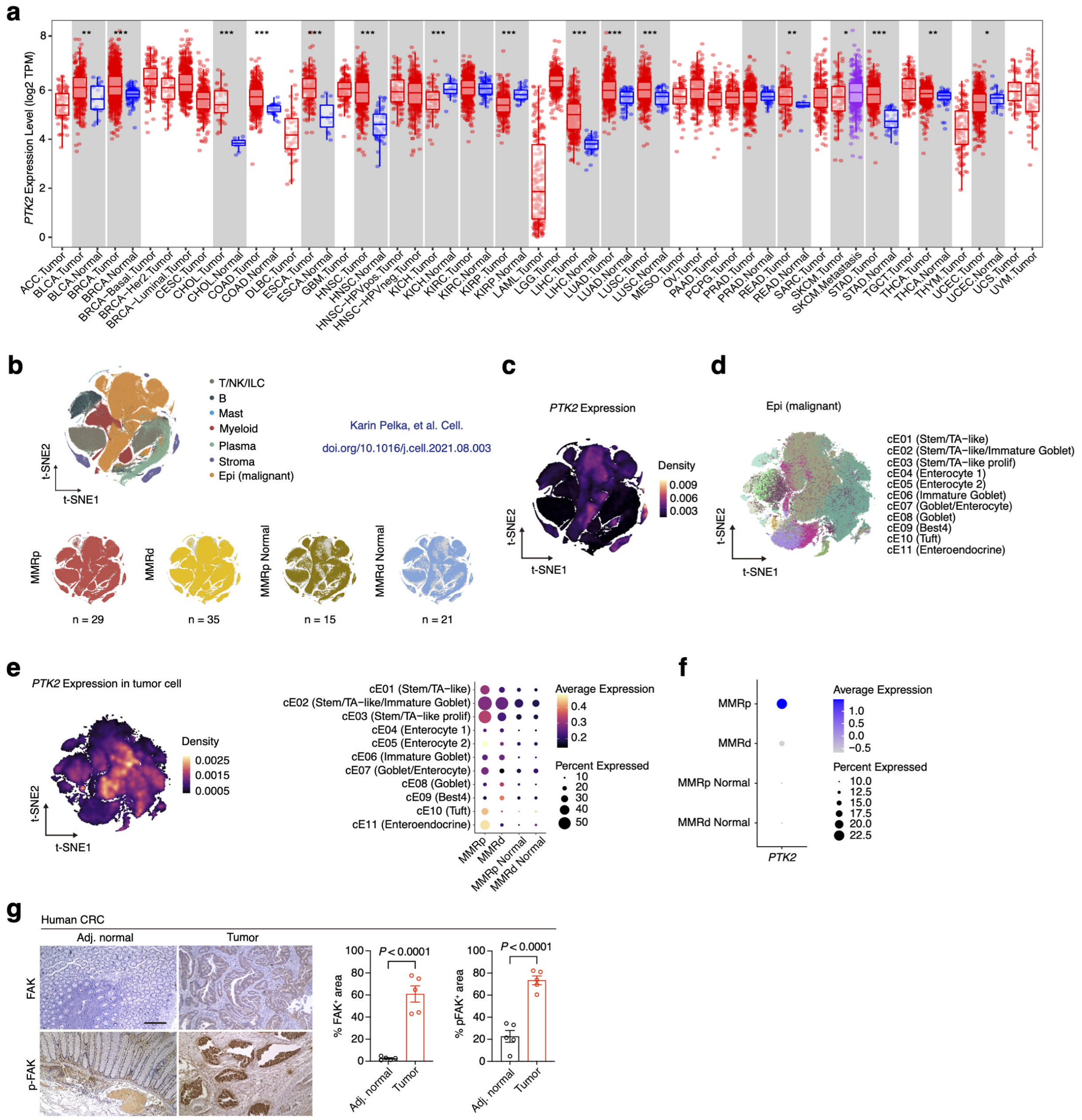

**Supporting Data Fig. 1**

**FAK is hyperactivated in human CRC**

**a,** Expression of *PTK2* across cancers based on the Tumor Immune Estimation Resource version 2. **b**, t-SNEs (t-distributed stochastic neighbor embedding) by major cell partitions of all samples (left). MMRp tumors (n = 35), MMRd tumors (n = 29), and MMRp and MMRd normal (n = 36) from 62 individuals (34 MMRd and 28 MMRp). **c**, t-SNE plot of all cells shown in **Supporting Data Fig. 1b**, with *PTK2*-expressing cells colored according to the cell density. **d**, t-SNE plot of tumor cells in the scRNA-seq data clustered into 11 subtypes. **e,** t-SNE plot of tumor cells shown in **Supporting Data Fig. 1d**, with *PTK2*-expressing cells colored according to cell density (left). Expression of *PTK2* across 11 tumor cell subtypes, where the dot size represents the proportion of PTK2-expressing cells and color indicates the average gene expression level (right). **f**, *PTK2* expression in the MMRp, MMRd, MMRP and MMRd normal groups. The dot size indicates the proportion of expressing cells, and the color represents the average gene expression level. **g**, Representative immunohistochemistry images of total and phosphorylated FAK (p-FAK, Tyr397) in human adjacent normal colon and CRC tumor tissues. The data are presented as the means ± SEMs (n = 5), as determined by two-tailed unpaired t tests.

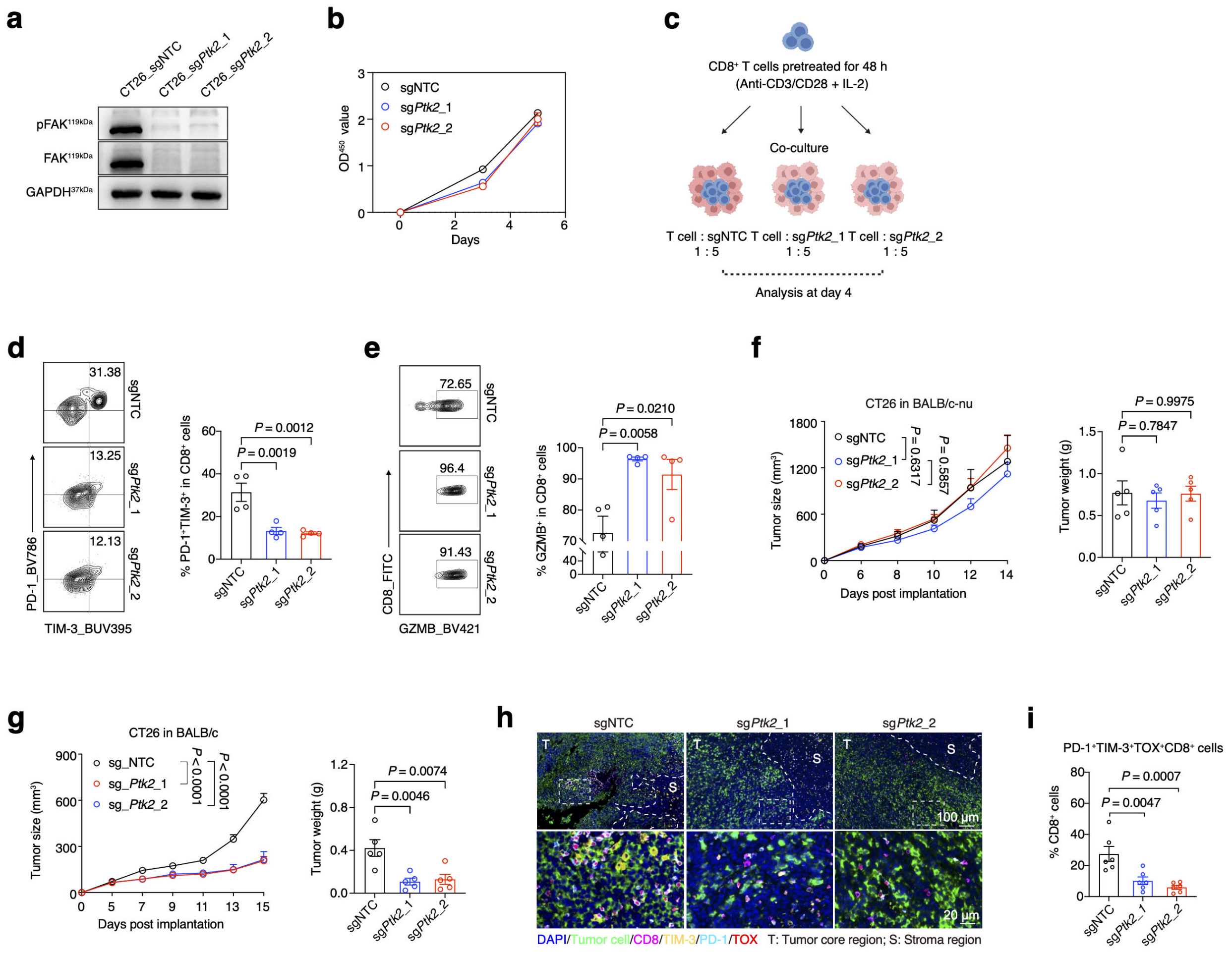

**Extended Data Fig. 1**

**Tumor cell-intrinsic FAK knockout reduces T-cell exhaustion in the CRC TME**

**a**, Western blot analysis of FAK/pFAK expression in CT26_sgNTC and CT26_sg*PTK2* cell lysates. **b**, Cell viability measurements of CT26_sgNTC and CT26_sg*PTK2* cells at days 3 and 5. **c**, Schematic diagram illustrating the *in vitro* coculture system of $CD8^+$ T cells with CT26_sgNTC and CT26_sg*PTK2* tumor cells. **d**, Percentages of $PD\text{-}1^+TIM\text{-}3^+CD8^+$ T cells among $CD8^+$ T cells in coculture systems (n = 4 per group). The data are presented as the means ± SEMs, as determined by one-way ANOVA. **e**, Percentage of $GZMB^+CD8^+$ T cells among $CD8^+$ T cells in the coculture system (n = 4 per group). The data are presented as the means ± SEMs, as determined by one-way

ANOVA. **f**, Tumor growth curves and tumor weights of CT26_sgNTC and CT26_sg*PTK2* tumors in BALB/c-nude mice. The data are presented as the means ± SEMs (n = 5 mice per group), as determined by two-way ANOVA or one-way ANOVA. **g**, Tumor growth curves and weights of CT26_sgNTC and CT26_sg*PTK2* tumors in BALB/c mice. The data are presented as the means ± SEMs (n = 5 mice per group), as determined by two-way ANOVA or one-way ANOVA. **h** and **i**, Representative mIHC images of CT26_sgNTC and CT26_sgPTK2 tumors (blue), tumor cells (green), CD8+ T cells (pink), PD-1+ T cells (cyan) and TOX+ T cells (red) (left). Comparisons of the percentages of terminally exhausted T cells (PD-$1^{+}$TIM-$3^{+}$$TOX^{+}$) among $CD8^{+}$ T cells within tumor core regions from CT26_sgNTC and CT26_sg*PTK2* tumors. The data are presented as the means ± s.e.m.ss (n = 6 ROIs per group), as determined by one-way ANOVA.

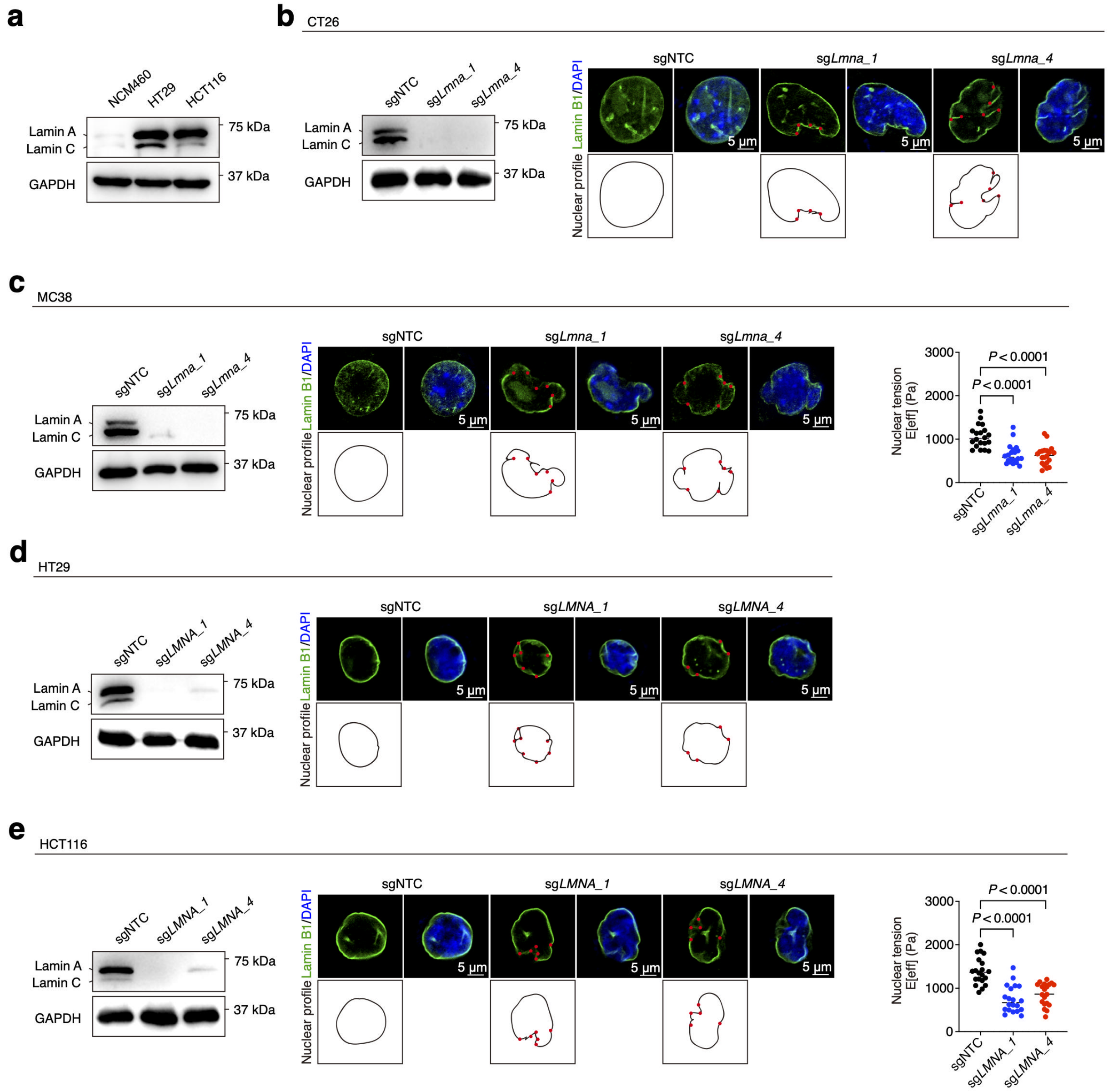


### Extended Data Fig. 2

### Lamin A/C is essential for the maintenance of nuclear tension

**a**, Western blot analysis of Lamin A/C expression in NCM460, HT29 and HCT116 cell lysates. **b**, Western blot of Lamin A/C expression in CT26_sgNTC and sg*Lmna* cell lysates (left). Images of the Lamin B1-labeled nucleus and nuclear profile in CT26_sgNTC and sg*Lmna* cells. The red points indicate the nuclear folds (right). **c**, Western blot of Lamin A/C expression in MC38_sgNTC and sg*Lmna* cell lysates (left). Images of the Lamin B1-labeled nucleus and nuclear profile in MC38_sgNTC and sg*Lmna* cells. The red points indicate the nuclear folds (middle). Measurements of the nuclear tension of MC38_sgNTC and sg*Lmna* cells (right, n = 20). **d**, Western blot of Lamin A/C expression in HT29_sgNTC and sg*LMNA* cell lysates (left). Images of the Lamin B1-labeled nucleus and nuclear profile in HT29_sgNTC and sg*Lmna* cells. The red points indicate the nuclear folds (right). **e**, Western blot of Lamin A/C expression in HCT116_sgNTC and sg*LMNA* cell lysates (left). Images of the Lamin B1-labeled nucleus and nuclear profile in HCT116_sgNTC and sg*LMNA* cells. The red points indicate the nuclear folds (middle). Measurements of the nuclear tension of HCT116_sgNTC and sg*LMNA* cells (right, n = 20).

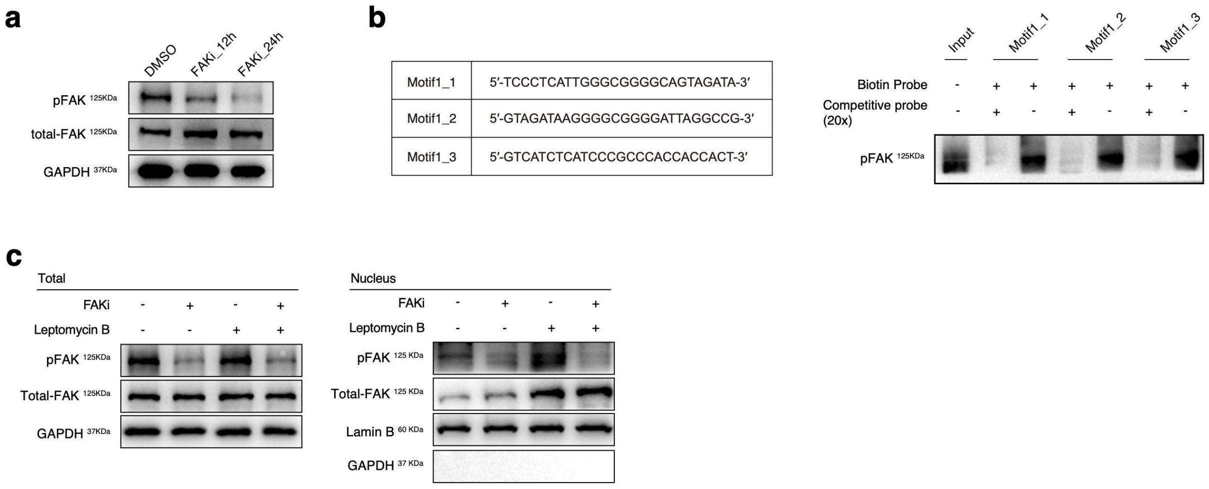


**Extended Data Fig. 3**

**Tumor nuclear pFAK promotes IL-6 secretion by regulating the transcription factor SP1**

**a**, Western blot analysis of total FAK and pFAK expression in CT26 cells treated with DMSO or FAKi for 12 and 24 h. **b**, DNA pull-down analysis showing the binding of pFAK to *Sp1* promoters at the predicted top motif. **c**, Western blot analysis of total and nuclear FAK/pFAK expression in FAKi (24 h)- and LMB (1 h)-treated CT26 cells.

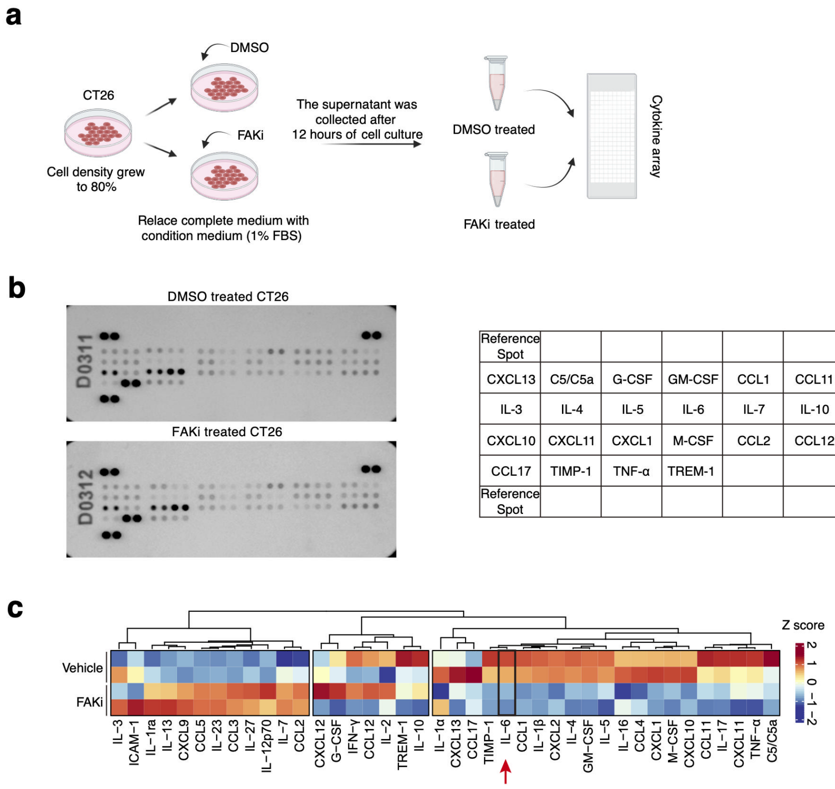


**Extended Data Fig. 4**

**pFAK regulates the secretion of several chemokines and cytokines, including IL-6, in CT26 tumor cells**

**a-b**, Cytokine assay of cytokines in the supernatants of DMSO- or FAKi-treated CT26 cells. **c**, Heatmap generated by hierarchical clustering of changes between DMSO- and FAKi-treated CT26 cells. The color and intensity indicate the relative protein expression levels in the culture medium.

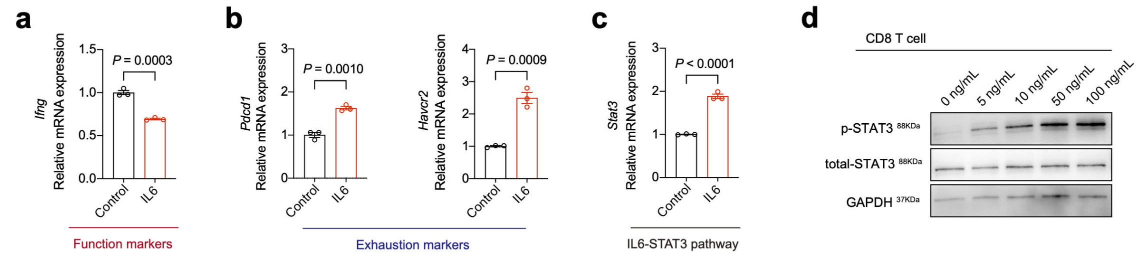


**Extended Data Fig. 5**

**IL-6 induces T-cell exhaustion**

**a–c,** Real-time quantitative PCR (RT–qPCR) analysis of *Ifng*, *Pdcd1, Havcr2,* and *Stat3* transcript levels in IL-6-treated CD8$^+$ T cells. CD8$^+$ T cells were treated with IL-6 (10 ng/mL) for 3 days (n = 3). Data are presented as the means ± SEMs, as determined by two-tailed unpaired t tests. **d**, Western blot analysis of pSTAT3 and total STAT3 expression in CD8$^+$ T cells treated with a dose gradient of IL-6. The dose-dependent experiment was performed with a fixed IL-6 treatment time of 30 min.

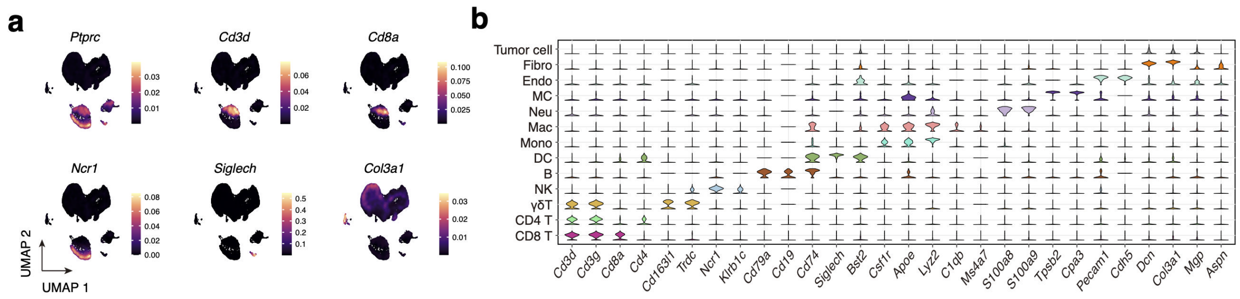


**Extended Data Fig. 6**

**Cell subpopulation markers for scRNA-seq data from vehicle- and FAKi-treated tumors**

**a,** Expression density distribution of known cell type marker genes on the UMAP plot from **Fig. 5e**, where brighter colors indicate higher concentrations of cells expressing the gene. **b,** Violin map showing the expression levels of specific markers in each cell type.

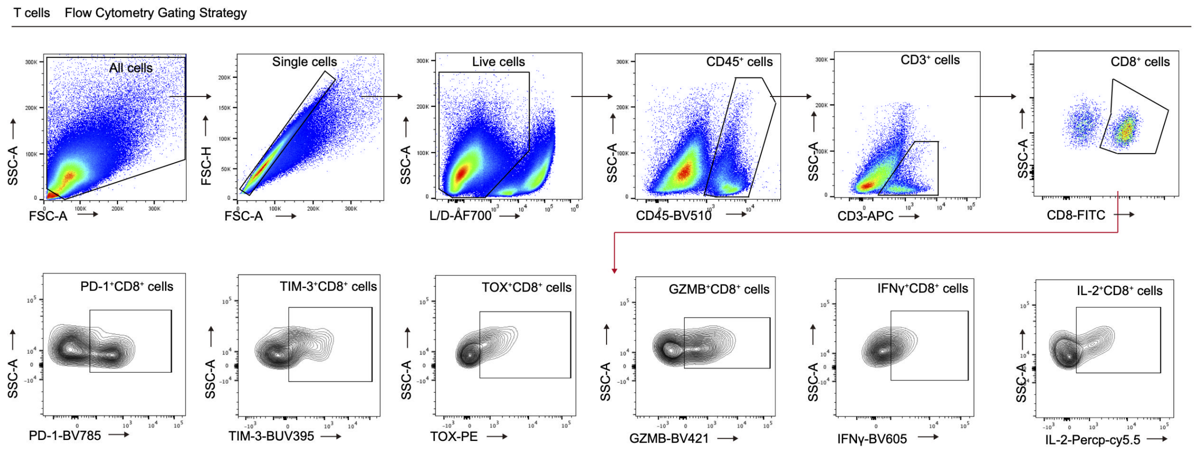


**Extended Data Fig. 7**

**Flow cytometry gating strategy for T cells**

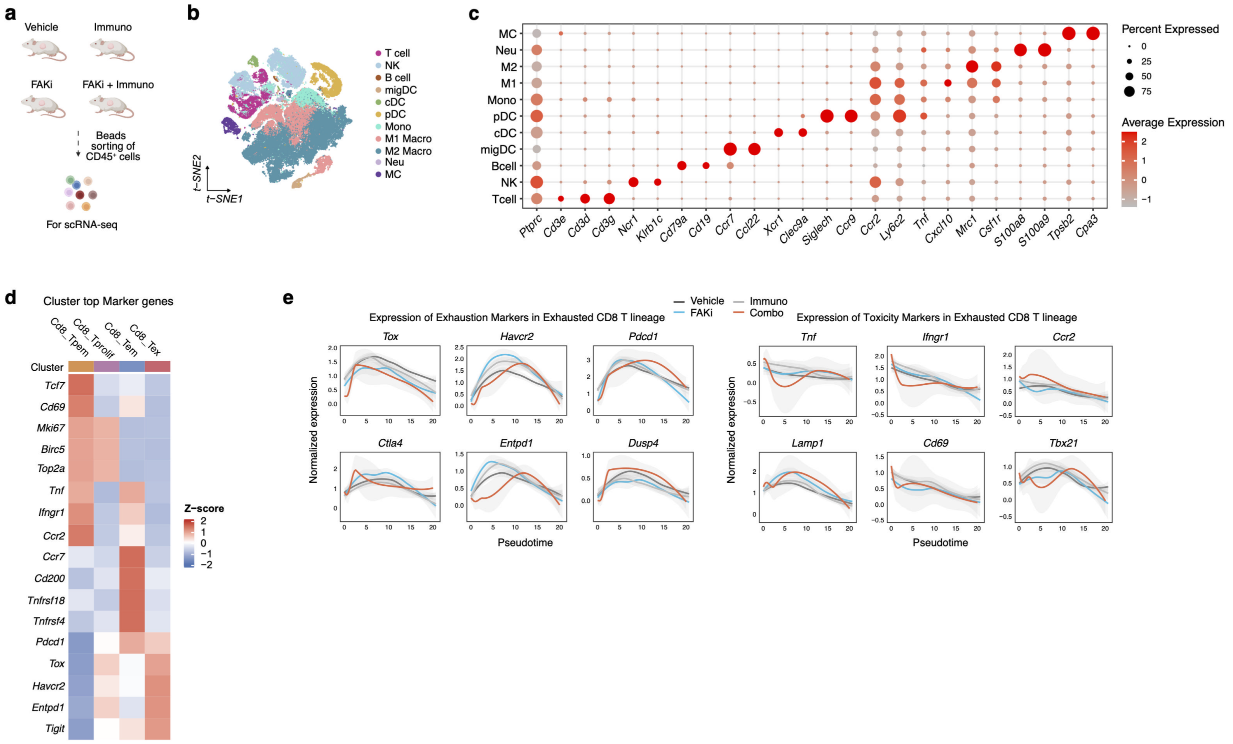


### Extended Data Fig. 8

**Cell subpopulation markers for scRNA-seq data from vehicle-, immune-, FAKi- and combination-treated tumors**

**a,** Schematics depicting the $CD45^{+}$ cell sorting strategy for subsequent scRNA-seq. **b**, t-SNE plot of 75,631 cells from vehicle-, immuno-, FAKi- or combination-treated tumor tissues (n = 2 per group), colored according to the annotated cell type. **c**, Dot plot showing the expression levels of specific cell type markers in each cell type. The dot size indicates the proportion of expressing cells, and the color represents the average gene expression level. **d**, Heatmap showing the expression levels of specific markers in the $CD8^{+}$ T-cell subtype identified in **Fig. 6f**. **e,** Plot of exhaustion and cytotoxicity markers along the exhausted $CD8^{+}$ T-cell lineage based on pseudotime from **Fig. 6i**. The black line represents the vehicle group, the gray line represents the immunotherapy group, the blue line represents the FAKi group, and the red line represents the combination-treated group.
